# Exploring eco-evolutionary and temporal patterns of arbuscular mycorrhizal fungal communities colonizing *Sorghum bicolor* across sites of contrasting land use history and climate

**DOI:** 10.1101/2025.10.07.681064

**Authors:** Philip A. Brailey-Crane, Sedona Marie Spann, Thomas H. Pendergast, Ben J. Long, Ashton K. Brinkley, Sarah R. Mondibrown, Nancy Collins Johnson, Katrien M. Devos, Jeffrey L. Bennetzen

## Abstract

**Societal Impact Statement:** *Sorghum bicolor* is a globally important crop, having endpoint uses ranging from human food to animal feed to biofuel production. Sorghum is stress-tolerant and can be grown on marginal land that may be otherwise unsuitable for large-scale food production. Sorghum is therefore a promising candidate for agricultural strategies focused on maximizing production on these marginal lands through beneficial interactions with arbuscular mycorrhizal fungi and other microbes. This study investigates how sorghum genotypes and their interactions with the environment can be leveraged to foster particular AMF assemblages which can be investigated further for their effect on the host plant.

Arbuscular mycorrhizal fungal (AMF) symbiosis can influence crop production, but can be variable across environmental conditions, host-partner complementarity and temporal dynamics. Understanding how these factors interact to shape AMF community assembly allows for the selection of crop genotypes that may maximally utilize AMF associations in agricultural systems. We assessed the development of AMF communities colonizing the roots of eight genetically diverse genotypes of *Sorghum bicolor* across a growing season. We used two field sites with contrasting environments and management histories. Sorghum cultivated in Arizona (AZ) contained low diversity AMF communities, while in Georgia (GA) sorghum harbored more diverse and evenly distributed AMF communities. We observed evidence of host-filtering of AMF communities, though with genotypes displaying more distinct associations in GA than AZ. AZ showed rapid shifts from early *Funneliformis mosseae* dominance to dominance by either *Entrophospora etunicata* or *Diversispora aurantia*. In GA, such drastic abundance shifts were not observed. Instead, consistent temporal turnover was associated more with higher level family abundance patterns driven by the combination of minor variations in multiple low-abundance taxa. Our findings demonstrate that there is potential for leveraging intra-species genetic variation in AMF community assembly as an extended plant phenotype.

## Introduction

Arbuscular mycorrhizal fungi (AMF) are obligate plant symbionts which form associations with over 70% of land plants (Brundrett, 2009; van der Heijden et al., 2015), including many commercial crops, such as *Sorghum bicolor*. AMF can facilitate nutrient uptake (Fitter et al., 2011; Hodge et al., 2010; Neumann & George, 2010) and improve biotic and abiotic stress resistance of the host (Yang et al., 2014), and contribute to soil health maintenance (Jeffries et al., 2003). AMF are prevalent across agroecosystems (Oehl et al., 2017), though their abundance and functional contributions vary with agronomic practices (Manoharan et al., 2017; Verbruggen et al., 2015). Host crops also display a wide range of intra-species capacities to form and benefit from AMF associations (De Vita et al., 2018), and in modulating AMF community assembly (García de León et al., 2020). Modern crop cultivars have mostly been developed for high input agricultural systems, with altered plant physiological traits that engender downstream consequences on mycorrhizal capacity (Hetrick et al., 1993; Londoño et al., 2019; Wang et al., 2023). Plant phenotypes linked to AMF capacity can include root exudation patterns (Steinkellner et al., 2007), root architecture (Cockerton et al., 2020; Sweeney et al., 2021), and overall physiology and nutrient use efficiency (Cockerton et al., 2020), which can also be highly plastic depending on environmental conditions (Wen et al., 2022).

There is a growing consensus that natural soil microbial diversity in agricultural systems could be utilized to more sustainably provide crop yields under reduced input systems (Brito et al., 2021; Gianinazzi et al., 2010; Thirkell et al., 2017). This is particularly relevant to sorghum, a globally important crop capable of yielding on marginal lands where other traditional agricultural systems may not be productive (Hazmi et al., 2022; Mar et al., 2019). In a US context, sorghum is a key strategy component to generating renewable fuel sources with low environmental impacts (Dahlberg, 2019; Morrissey et al., 2021; Rao et al., 2015). Optimizing sorghum production on marginal land would provide a suitable case study for harnessing the natural biological capital of soils (Regassa & Wortmann, 2014). Numerous genetic resources have been developed to facilitate sorghum breeding (Boatwright et al., 2022; Brenton et al., 2016; Cuevas et al., 2023). These resources have been so far utilized mostly for aboveground agrological trait analyses but could be leveraged to assess the diversity and benefits of AMF associations.

Examining AMF communities at only one point during crop development to determine Genotype x Environment (GxE) interactions can inadvertently mask key dynamics, as AMF also display strong temporal dynamics across spatial scales in both abundance and community assembly patterns. Within a single root system, colonizing AMF exhibit high turnover within the lifespan of a plant (Dang et al., 2021; Gao et al., 2019; Lu et al., 2019). More broadly across whole ecosystems, AMF communities also demonstrate turnover tracking seasonal shifts in the environment, and in interaction with aboveground plant community successional patterns both in natural communities and agricultural rotations across seasonal and longer time scales (Bainard et al., 2014; Dumbrell et al., 2011; Oehl et al., 2011). In the context of a single plant life-cycle, this may be attributed to a wide phenotypic range of life history traits found in AMF, including biomass allocation patterns (Weber et al., 2019), rates of growth, longevity and competitive ability against other fungi (Hart & Reader, 2002; Hart et al., 2001; López-García et al., 2014). Such changes in AMF community compositions and/or roles may also be associated with host-plant shifts in resource allocation during development (Abrar et al., 2024; Buade et al., 2021), though truly untangling plant growth stage from temporal shifts in abiotic conditions in field systems is difficult.

In this study, we investigated how sorghum genotype and environment interact to shape AMF community composition and temporal dynamics. We grew eight genetically diverse sorghum genotypes from the sorghum Bioenergy Association Panel (BAP, Brenton et al., 2016) at two ecologically distinct field sites in Georgia (GA) and Arizona (AZ), USA. These sites represent contrasting climates and soil conditions, allowing insight into host genotype differentiation across sites. We also tracked AMF community development across a growing season, sampling roots from 22 to 129 days after planting to capture temporal progression of community assembly. We hypothesized that (1) different sorghum genotypes would associate with distinct AMF communities, reflecting genotype-specific compatibility or co-selection, (2) the environmental context would mediate these associations through processes such as acting as a filter of the available AMF community or through changing plant resource needs from AMF, and (3) that AMF communities would show strong temporal shifts that would also differ across sites. This work provides a multi-dimensional view of plant-AMF interactions under field conditions, highlighting the potential for monitoring these dynamics to select and breed crops with enhanced compatibility with beneficial AMF.

## Methods

### Experimental Field Design and Sampling

Root samples were collected from paired field experiments conducted at the Wellbrook site of the J. Phil Campbell Research and Education Center, GA, USA (33.89632°, −83.422129°) and Maricopa Agricultural Center, AZ USA (33.06975°, −111.97901°), between April and September 2021. The two sites are located within contrasting climates / biomes (AZ: Köppen BWh, hot desert climate, Sonoran Desert biome; GA: Köppen Cfa, humid subtropical climate, temperate broadleaf and mixed forest biome) (Supplementary Figure S1), and having contrasting soil edaphic properties (Table 1) and management histories. In September 2018, the GA site was treated with glyphosate at a rate of 2.34 L ha^-1^ and sowed with ryegrass (*Lolium multiflorum, cv.* Marshall) the following month. Nitrogen was applied at 56.1 kg ha^-1^ as a 32% solution of urea-ammonium nitrate in April 2019 but otherwise the field was left fallow until our experiment. To prepare the plot in GA, the experimental area was again treated with glyphosate at a rate of 4.68 L ha^-1^ on April 27^th^, 2021, and limed at a rate of 3362.86 kg ha^-1^ on May 14^th^, 2021. The AZ field site had been bare soil fallow in 2017/2018, sown with barley (*Hordeum vulgare*) in 2019, and bare soil fallow again in 2020/21 prior to being sown with sorghum for our trial. No herbicide or fertilizer treatments were applied to the AZ field site.

**Table 1.** Soil edaphic properties recorded at each field site. Soils were collected in 2022 (GA) and 2023 (AZ) from follow-up experiments conducted within the same field sites, using soils collected across three discrete experimental blocks (n=3 per block) to best capture a representative value for the field. Significant differences in properties between sites were determined through ANOVA applied to linear models (variable ∼ site). Replication: AZ n = 9; GA n = 9.

| Soil Edaphic Property | AZ | GA | F-value | P-value |
| --- | --- | --- | --- | --- |
| pH | 8.27 ± 0.04 | 6.35 ± 0.10 | 245.47 | 3.97 x 10<sup>-11</sup> *** |
| Ca (mg kg<sup>-1</sup>) | 6529.20 ± 196.74 | 667.67 ± 76.77 | 783.31 | 5.11 x 10<sup>-15</sup> *** |
| K (mg kg<sup>-1</sup>) | 149.04 ± 5.66 | 140.93 ± 17.35 | 0.42 | 0.53 |
| Mg (mg kg<sup>-1</sup>) | 278.07 ± 5.41 | 113.63 ± 9.74 | 151.76 | 1.41x10<sup>-9</sup> *** |
| Mn (mg kg<sup>-1</sup>) | 9.68 ± 0.71 | 17.86 ± 4.18 | 2.50 | 0.13 |
| P (mg kg<sup>-1</sup>) | 282.05 ± 6.14 | 4.86 ± 0.86 | 2458.71 | 6.01x10<sup>-19</sup> *** |
| Zn (mg kg<sup>-1</sup>) | 0.57 ± 0.07 | 0.91 ± 0.19 | 2.09 | 0.17 |
| NO<sub>3</sub>-N (mg kg<sup>-1</sup>) | 3.86 ± 0.45 | 4.20 ± 0.57 | 0.13 | 0.72 |
| NH<sub>4</sub>-N (mg kg<sup>-1</sup>) | 1.29 ± 0.05 | 7.25 ± 0.76 | 103.78 | 2.12x10<sup>-8</sup> *** |
| LOI Organic Matter (%) | 0.91 ± 0.02 | 4.74 ± 0.44 | 133.67 | 3.52x10<sup>-9</sup> *** |

Eight sorghum genotypes, one from each of eight subpopulations (Brenton et al., 2016), were selected from the Bioenergy Association Panel (BAP (Brenton et al., 2016) to capture a wide genetic diversity (Supplementary Table S1). (Brenton et al., 2016) defines six clusters very broadly aligned with botanical race, though we chose to consider eight potential genetic sub-populations following the Evanno method (Supplementary Figure S2). Genotypes were obtained from the USDA-ARS Germplasm Resources Information Network (GRIN, https://www.ars-grin.gov).

Sorghum was sowed in accordance with planting schedules typically used at the two locations (Table 2). Three replicate plots for each genotype were planted at both field sites. Individual genotype plots were organized in linear rows of 18 plants spaced approximately 20 cm apart. Distance between genotypic plots within a row was ∼140 cm, and inter-row distance was ∼90 cm. Genotype plots were distributed randomly within replicate blocks (Supplementary Figure S1).Two seeds were planted at each position and thinned to one plant after emergence. In GA, there was low seedling success, resulting in some plants removed during thinning being transferred to another replicate plot where appropriate. Plots in GA were hand-weeded every few days for the first eight weeks of the study, and approximately weekly for the remainder of the experiment. Plots in AZ were hand weeded approximately every two weeks, at the same time as experimental sampling. The GA field site was irrigated as needed during low rainfall periods. The AZ field site was flood irrigated approximately every two weeks.

**Table 2.** Timetable for planting and root collection in GA and AZ. Timepoint designations are unified by similar plant ages.

| Unified Timepoint (TP) | AZ |  | GA |  |
| --- | --- | --- | --- | --- |
|  | Date | Plant Age<sup>1</sup> (dap) | Date | Plant Age<sup>1</sup> (dap) |
| Sowing | 28<sup>th</sup> Apr 21 | - | 2<sup>nd</sup> Jun 2021 | - |
| TP1 | 19<sup>th</sup>-21<sup>st</sup> May 21 | 22-24 | *No collection* |  |
| TP2 | 1<sup>st</sup>-2<sup>nd</sup> Jun 21 | 35-36 | *No collection* |  |
| TP3 | 23<sup>rd</sup>-24<sup>th</sup> Jun 21 | 57-58 | 22<sup>nd</sup>-23<sup>rd</sup> Jul 2021 | 51 |
| TP4 | 12<sup>th</sup>-13<sup>th</sup> Jul 21 | 76-77 | 9<sup>th</sup> -10<sup>th</sup> Aug 21 | 69 |
| TP5 | 26<sup>th</sup>-27<sup>th</sup> Jul 21 | 90-91 | 25<sup>th</sup>-26<sup>th</sup> Aug 21 | 85 |
| TP6 | 9<sup>th</sup> -10<sup>th</sup> Aug 21 | 104-105 | 8<sup>th</sup>-9<sup>th</sup> Sep 21 | 99 |
| TP7 | 2<sup>nd</sup>-3<sup>rd</sup> Sep 21 | 128-129 | 8<sup>th</sup> Oct 21 | 129 |

<sup>1</sup> Plant age = days after planting (dap). For analysis, the earliest dap at each TP is used when calculating temporal distances.

For between-site comparisons, sampling across the two sites was unified by bandings of plant age at harvest classified as categorical timepoints. For within-site temporal assessments, the actual plant ages (days after planting, dap) were used (**Table 2**) to accommodate models that can test non-linear associations over time. Where possible, two plants were collected per replicate plot at each time point and pooled for analysis. In GA, roots were washed at the field site to remove soil and debris, and kept on ice until storage at −20°C. In AZ, the collected roots were stored at 4°C until washing and then stored at –20°C within 48 hours. Sorghum dry biomass and height was recorded at the final harvest for both sites (TP7).

### DNA Extraction and Library Preparation

Washed roots were ground in liquid nitrogen with a mortar and pestle. DNA was extracted from ∼300 mg of powdered roots using a CTAB protocol adapted from (Xin & Chen, 2012). Briefly, samples were mechanically disrupted, and DNA was phase-separated from other components through CTAB/phenol-chloroform addition. Nucleic acids were precipitated in ethanol and solubilized in 10 mM Tris-HCl (pH 8.0), and DNA concentration recorded by spectrophotometry (ND-1000, Nanodrop). Where two plants were collected per replicate, the extracts were pooled in equal amounts prior to PCR amplification.

To assess AMF community composition, we generated amplicon libraries from the ribosomal large subunit (LSU) gene region. This was chosen over the more commonly adopted small subunit (SSU) region due to the limited application of SSU amplicons to reflect known defined species level differences in comparison to the LSU region (Schlaeppi et al., 2016). There are also increasing amounts of phylogenetic and bioinformatics resources available for the LSU region (Delavaux et al., 2022, 2024, 2021; Tedersoo, et al., 2024). FLR3 (TTGAAAGGGAAACGATTGAAGT, Gollotte et al., 2004) and FLR2 (GTCGTTTAAAGCCATTACGTC, van Tuinen et al., 1998) were chosen as the forward and reverse primers, respectively, generating target amplicons of approximately 300-400 bp, a length that can be fully sequenced by Illumina sequencing technologies without needing to concatenate non-overlapping forward and reverse reads (per (Delavaux et al., 2022)). The chosen primer pair resulted in some non-target amplification of non-AMF sequences of a shorter length than expected.

Barcoded amplicon libraries were generated using a two-step PCR approach based on (Kozich et al., 2013). Initial PCRs were carried out in 8 µl reaction volumes containing 2 µl of 5ng/µl DNA input, 10nM each of FLR3 and FLR2 primers, and a final concentration of 1X Phusion Hot start master mix (Thermo Fisher Scientific). PCR conditions were 95°C 5 min initial denaturation, 35 cycles: 94 30s, 55 30s, 72 30s, followed by 72° 5 min final elongation. FLR3 and FLR2 primers were modified to contain 12 bp spacer sequences (NNNNNHNNNNNN) to create random variation to aid in cluster delineation during Illumina sequencing, and regions complementary to the Illumina indexing primer for the step 2 barcode PCR. Initial PCR reactions were diluted 1:15 with 10 mM Tris-HCl (pH 8.0) for input into Step-2 PCR, which was carried out in 15 µl reaction volumes containing 2 µl of diluted Step-1 PCR product (PCR conditions: 95°C 3 min initial denaturation, 10 cycles: 95°C 30s, 56°C 30s, 72°C 30s, followed by 72°C 5 min final elongation). Barcoded amplicon libraries were pooled in equimolar amounts for sequencing. AMF detection was low from this amplicon, and sequencing was performed twice for some samples due to non-specific amplification of amplicons in the ∼220-300 bp size class through this protocol that are preferentially sequenced over longer amplicons by Illumina technologies. A further subset of libraries did not yield enough sequences attributable to AMF through two sequencing runs and so PCR was performed again for these samples using the same cycling conditions in 25 µl reactions containing 4 µl of non-normalized DNA input. For this third sample set, bands of the expected size class were excised from 2% agarose gels and diffused from the gel in 500 µl of 10 mM Tris-HCl (pH 8.0) for 30 minutes to be used as input for the Step-2 barcode PCR reactions following the same protocol previously stated for the Step-2 PCR. Amplicon libraries were sequenced across three Illumina NextSeq 2000 P2 flow cell runs (2 x 300 bp reads). AMF community data, including alpha diversity values, ASV abundances and AMF read counts from each run can be found in Supplementary File S2.

### Sequence Data Processing and Phylogeny-Aware Assignment of Taxonomy

Primers were trimmed from the forward and reverse reads using *Cutadapt v4.5* (Martin, 2011). To accommodate sequences shorter than 300 bp, sequences were also trimmed for the reverse complement of the appropriate primer sequences to remove through-sequencing of adapters. Trimmed sequences were quality-checked using *fastQC* (http://www.bioinformatics.babraham.ac.uk/projects/fastqc/). Amplicon sequence variants (ASVs) were determined using the *dada2 v1.28.0 R* package (Callahan et al., 2016). Sequences were further trimmed to 230bp or discarded if smaller, then quality filtered, dereplicated, denoised, and chimera filtered following the standard dada2 pipeline (Callahan et al., 2016a; Callahan et al., 2016b), with the exception of performing denoising in pooled mode. Additional *de novo* chimera filtering was carried out in QIIME2 followed by a dataset-wide abundance filtering (absolute count > 10 across the entire dataset and present in > 13 samples).

Adapting the protocol of (Delavaux et al., 2022) for AMF phylogeny-informed taxonomy assignments, ASV sequences were subjected to a BLAST screening to remove both non-homologous sequences and homologous sequences with query coverage ≤ 80% against the top hit using a Glomeromycota-only subset of the EUKARYOME database v1.9.3 (Tedersoo, Hosseyni Moghaddam, et al., 2024). Taxonomic designations were assigned to the screened ASVs using a naïve Bayes classifier trained on a fungal-only subset of the EUKARYOME database implemented in QIIME2 v2024.2 (Bolyen et al., 2019). We chose to use a broad fungal reference dataset to ensure that no sequences were spuriously assigned to AMF through this method due to a lack of alternative possibilities. ASVs that could be assigned to the phylum *Glomeromycota* were aligned against the Delavaux backbone phylogeny *v.16 2024* (Delavaux et al., 2024) using *mafft v7.520*, (Katoh & Standley, 2013) with the E-INS-i parameters. Phylogenetic trees were constructed using *RAxML v8.2.12* (Stamatakis, 2014). Maximum likelihood / rapid bootstrap trees (n=500) were constructed using the GTRGAMMAI algorithm, followed by the construction of a majority rule (>50% bootstrap replicates) consensus tree. The phylogeny was visualized and annotated using the interactive Tree of Life (iTol, (Letunic & Bork, 2024). Where unassigned ASVs sat within clades of assigned ASVS, their assignments were updated to match their contemporaries. Where unassigned ASVs did not sit clearly within existing defined clades we considered both bootstrap support (>50%) and visual branch length differences to hypothesize novel molecular taxa (MT). Phylogenetic annotations and descriptions are expanded upon in Supplementary Figure S3, Supplementary File S2 and S3.

### Microbiome Parameters and Statistical Analyses

Unless stated, base figure generation and statistical analyses were performed using packages available within the *R* environment *v.4.3.1* as implemented in *Rstudio v.2023.6.1*, with some additional manual text-only annotations made. Three datasets were used for statistical analyses. To avoid the potential for bias of community succession patterns in between-site comparisons where non-overlapping categorical timepoints were sampled, a subset of the data was used (*Dataset GxE*) containing only samples from the timepoint collections taken at both sites (TP3-TP7, omitting TP1 and TP2, which were exclusively sampled in AZ). ‘Dataset GxE’ was used to analyze the relative contribution of GxE interactions in determining AMF community assembly. Two additional subset datasets were curated containing all timepoints sampled for AZ and GA independently. Descriptions of overall taxa abundances for the AZ site pertain to TP3-7 unless otherwise states, as early colonization is not highly representative of the community composition. These two datasets were used to analyze temporal patterns of AMF community assembly individually for each site (‘Dataset GA’ and ‘Dataset AZ’). Unless stated, data were analyzed based on linear models or linear mixed-effect models, where appropriate, constructed with *the nmle v.3.1.162* package and tested by ANOVA. Post-hoc pairwise comparisons of estimated marginal means were performed using the *emmeans v.1.10.5* package. Comparisons were conducted within multi-class factors, based on t-statistics, with p-values adjusted for multiple testing using the Tukey method. Statistical analyses performed and their results can be found in Supplementary File S3 and are summarized in the corresponding figures rather than in-text.

Rarefaction curves were generated per-sample to assess the completeness of AMF sampling. As it is difficult to determine whether low abundance sequences such as singletons are due to the true presence of rare taxa or barcode hopping, we performed a final per-sample abundance filter removing ASVs with an abundance < 0.1% per sample (Nikodemova et al., 2023) before aggregating ASVs belonging to the same species or molecular taxa ID as the main basis for further analyses using *phyloseq v.1.44.0* (McMurdie & Holmes, 2013). Taxonomic alpha diversity (species richness and Pielou’s Index) was also calculated per sample using *phyloseq*. Faith’s PD was calculated using the *picante v.1.8.2* (Kembel et al., 2010) package to assess phylogenetic diversity. All alpha diversity metrics were calculated using rarefied data to reduce the impact of sequencing depth on comparisons, and where Pielou’s index was not calculable (Species richness = 1, Shannon’s H’=0), we amended this value to zero to represent a completely uneven community for statistical analysis and plotting. To better understand the potential ecological processes underpinning community assembly over time we calculated the standardized effect sizes (SES) of both mean pairwise distances (MPD) between taxa within each sample, and the nearest taxon index (NTI) of each taxon within each sample. These were calculated based on species-level relative abundances using the *picante v. 1.8.2* package.

Variation in AMF community composition between comparisons was tested by PERMANOVA performed on Bray-Curtis distance matrices using the *vegan v. 2.6.4* package (Oksanen et al., 2013) using Hellinger-transformed species abundances. Pairwise PERMANOVA was implemented through *pairwiseAdonis v.0.5.1* (Arbizu, 2020) with Benjamini-Hochberg (BH) p-value corrections to control for multiple testing. Through these pairwise comparisons genotypes were aggregated into higher level groupings of community similarity for further testing. Principal Co-ordinate Analysis (PCoA) was performed to visualize AMF communities, also using Bray-Curtis distances based on Hellinger-transformed species abundances. Additional Constrained Analysis of Principal Co-ordinates (CAP) was performed, constraining communities by non-aggregated host-genotype. The ten AMF species most strongly associated with between-sample variation in this constrained ordination were tested for differences between our defined genotype community groups / clusters through linear mixed effect models using species abundances subjected to center log ratio (clr) transformation using the *microbiome v1.22.0* package (Lahti et al., 2017). As we had *a priori* assumptions about the abundance of the ten taxa tested across our genotype groups (which included genotypes shared between groups that can mask global variation), we carried out pairwise post-hoc comparisons for taxa found to be significantly different in the global ANOVA test regardless of whether BH-corrected p-values maintained significance for the global test. We report both BH-corrected and uncorrected p-values for these analyses for transparency, and for the post-hoc test considered pairwise differences significant only if they were upheld after Tukey p-value corrections.

Individual taxa (> 0.5% site-wide) relative abundance shifts over time were identified through generalized additive mixed models (GAMMs) using the R package *mgcv v.1.8.42* (Wood, 2011) for timepoint comparisons (Datasets GA and AZ), and through linear mixed-effect models for between-genotype (Dataset GxE) and between-site comparisons (Dataset GxE). This provided a global p-value for non-linear patterns. Post-hoc tests were performed based on linear models to identify specific shifts between treatment groups using *emmeans*, again using Tukey p-value corrections. Aggregate test results from between-site comparisons were visualized using iTol to display how indicator status cascades through taxonomic hierarchy.

## Results

### Study-wide statistics and taxonomic annotations

A total of ∼*9.9* million AMF-assigned sequences were generated across 282 samples. We recorded 915 Glomeromycota-assigned ASVs across all samples covering **76** species-level clusters, of which **32** were novel molecular taxa determined through phylogeny (Supplementary Figure S3, 4). Species belonged to 22 genera across **12** families. Six genera (and two families) were represented by a single species definition (*Acaulosporacea*/*Acaulospora*, *Archaeosporaceae*/*Archaeospora, Desertispora*, *Fuscutata, Funneliformis*, and *Microkamienskia*). Total sequencing depths and obtained AMF sequence depths varied across the experiment covering 256 – 168,618 AMF sequences per sample (median = 20k, See Supplementary Data S1 for per-sample sequencing depths both pre- and post-AMF filtering). Rarefaction analysis revealed that most samples began to approach an asymptote of ASV/species detection (Supplementary Figure S4).

### Genotype x Environment interactions of AMF community assembly

Site primarily determines community composition (**Figure 1A/B**, 38.4% of between-sample variation). Host-plant genotype and genotype x environment (GxE) interaction effects were also significant predictors of AMF community composition (p < 0.05), with relatively minor contributions (Genotype: **2.7%**; GxE: **3.1%**). Diversity also was lower in AZ than GA (Figure 1C). Pielou’s Index had the least pronounced difference between the two sites. Though not considered in our GxE comparisons, early in colonization (TP1-TP2/22-36 dap), samples in AZ exhibit dominance (> 50% relative abundance occupancy by a single species) in all samples but one by *Funneliformis mosseae* that would impact this comparison of evenness. When considering the common timepoints (TP3-TP7), **79.4** % of AZ samples exhibited still > 50% dominance-mostly by *F. mosseae* (n=47/118 samples), *Entrophospora etunicata* (n=24), or *Diversispora aurantia* (n=10). In contrast, only **26.7** % of GA samples displayed this dominance characteristic, with no set of species displaying a particular prevalence. All ten of most abundant (experiment-wide) genera have contrasting relative abundances between sites (**Figure 1D**, **Figure 2**), along with *Pervetustus, Cetraspora, Microdominikia*, and *Desertispora* (**Figure 2**). We observed 25 species and one genus as unique to GA, and none were unique to AZ (**Figure 2**). We found that 20 additional species were differentially abundant between the two sites. Four were in higher abundance in AZ (*F. mosseae*, *E. etunicata*, *D. aurantia*, *Desertispora omaniana*) and 16 were in higher abundance in GA (**Figure 2)**.

**Figure 1.**
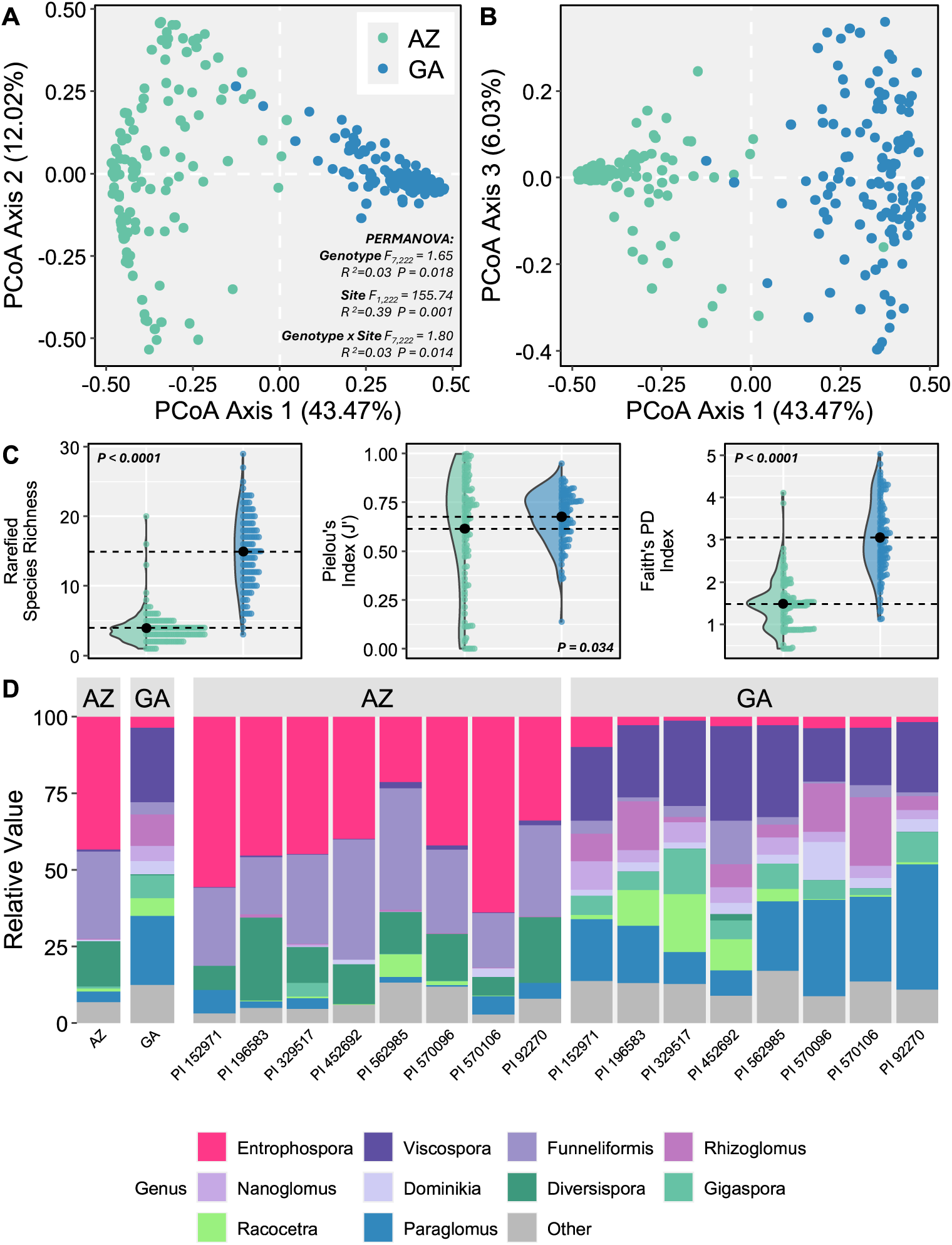
PCoA generated from Bray-Curtis distances between samples displaying **A)** Axis 1 and 2, **B)** Axis 1 and 3. P-values associated with PERMANOVA analysis performed comparing between genotypes, site and their GxE interaction are shown. **C)** Combined violin/bee swarm plots displaying the alpha diversity metrics of rarefied species richness, Pielou’s index of diversity (J’) and Faith’s phylogenetic diversity (PD) index based on species-level relative abundances at each site. Both raw and mean ± SE values are displayed. Only P-values associated with the main effect test of between-site differences for these panels. PCoA and alpha diversity plot samples are colored by site. Replication: AZ n=118; GA n=120. **D)** Taxa plots displaying mean relative abundances of genera, aggregated based on both site and genotype. Replication: AZ, n=14 for PI 196583 and PI 452692, n=15 for all other genotypes. GA, n=15 per genotype.

**Figure 2.**
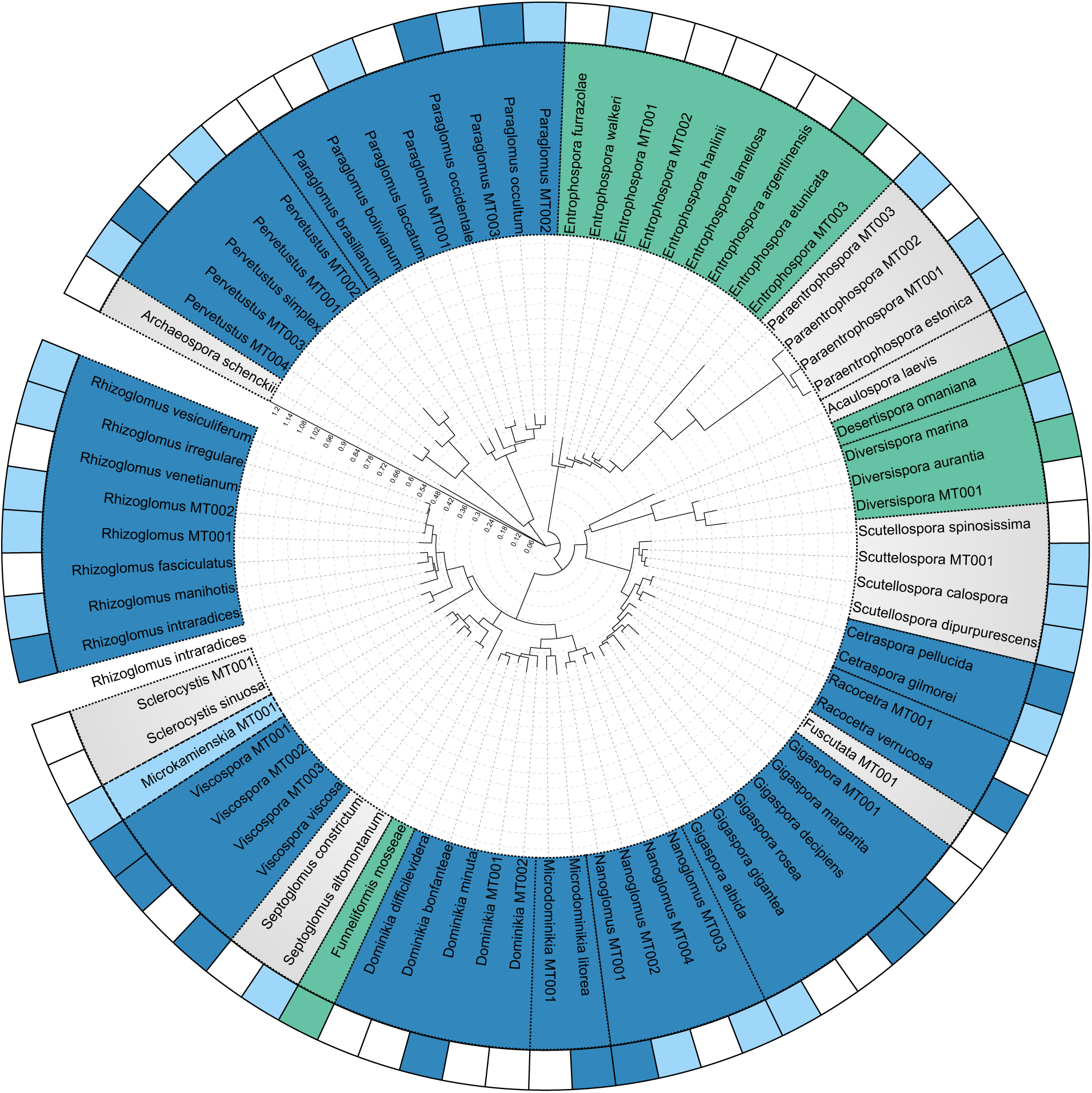
RAxML tree of AMF species found in this study, featuring one representative sequence per species / molecular taxa. Peripheral colored strips represent species and genera found to be significantly more abundant in either GA (blue) or AZ (green) or exclusively found in GA (light blue). The inner ring represents whether the aggregated genus was significantly more abundant in either site. Branch lengths represent phylogenetic distance between representative sequences, with the scale embedded within the tree.

The overall contribution of host genotype identity to AMF community composition is similar in GA than AZ, where it accounts for 9.99% and 9.57% of intra-site variation between samples respectively. In the CAP ordination, genotypes display clearer visual separation in GA however (Figure 4A/B). Pairwise-PERMANOVA analysis revealed that genotypes clustered into two groups in AZ and three groups in GA based on AMF community assembly, with some genotypes at each site being shared between the groupings (Supplementary Figure S5). Notably ‘group a’ identified in GA had no members that did not overlap membership with either ‘group b’ or ‘group c’. We focus our analyses on these emergent genotype groupings, which provide a clearer signal than direct genotype-by-genotype comparisons. In AZ, ‘group a’ contains three exclusive genotypes and ‘group b’ comprises a single exclusive genotype. Four genotypes exhibited joint membership to each group. In GA ‘group a’ comprises no exclusive genotypes, ‘group b’ comprises of three exclusive genotypes, and ‘group c’ comprises of one (Supplementary Figure S5). Genotypes cluster in different patterns at each site (Supplementary Figure S5), indicating that environmental filtering interacts with host-identity, as genotypes display different recruitment strategies at both sites in response to local climate/edaphic conditions and the available species pool. Notably, however, PI 570106 and PI 452692 harbor consistently divergent AMF communities from one another. This, in part, is related to *F. mosseae* abundance regardless of site, with PI 452692 being associated with a greater abundance of this taxon (**Figure 3 A/B**).

**Figure 3.**
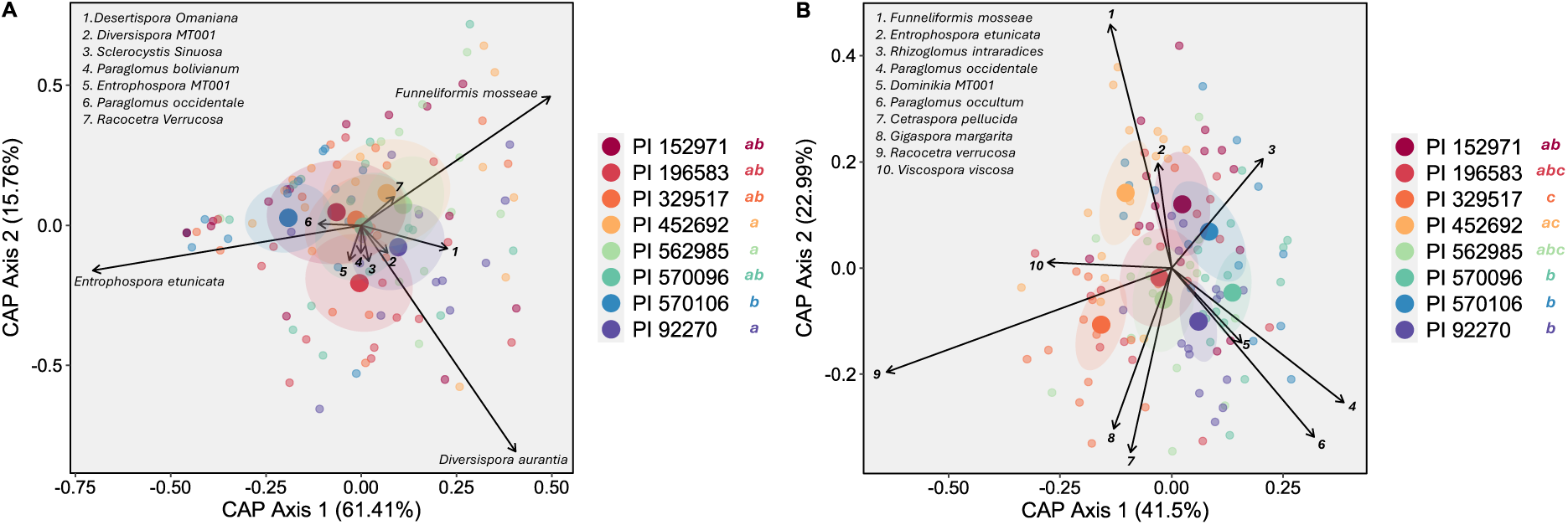
Canonical Analysis of Principal components (CAP) ordinations generated from Bray-Curtis distances between samples from dataset GxE. Constrained group centroids and constrained weighted averages of all samples are presented for **A)** AZ and **B)** GA. Ellipses around centroids display the standard error of the grouping variable. Arrow vectors display constrained species scores and represent the contribution of each species towards the separation of samples along the constrained axes 1 and 2. Only the ten most maximally informative species per site are included. Letters next to genotype labels represent groupings of statistical similarity (BH-adjusted p < 0.05) after pairwise PERMANOVA testing following global PERMANOVA model results presented in Figure 1. All samples are separated by genotype. Replication: AZ, n=14 for PI 196583 and PI 452692, n=15 for all other genotypes; GA, n=15 per genotype.

**Figure 4.**
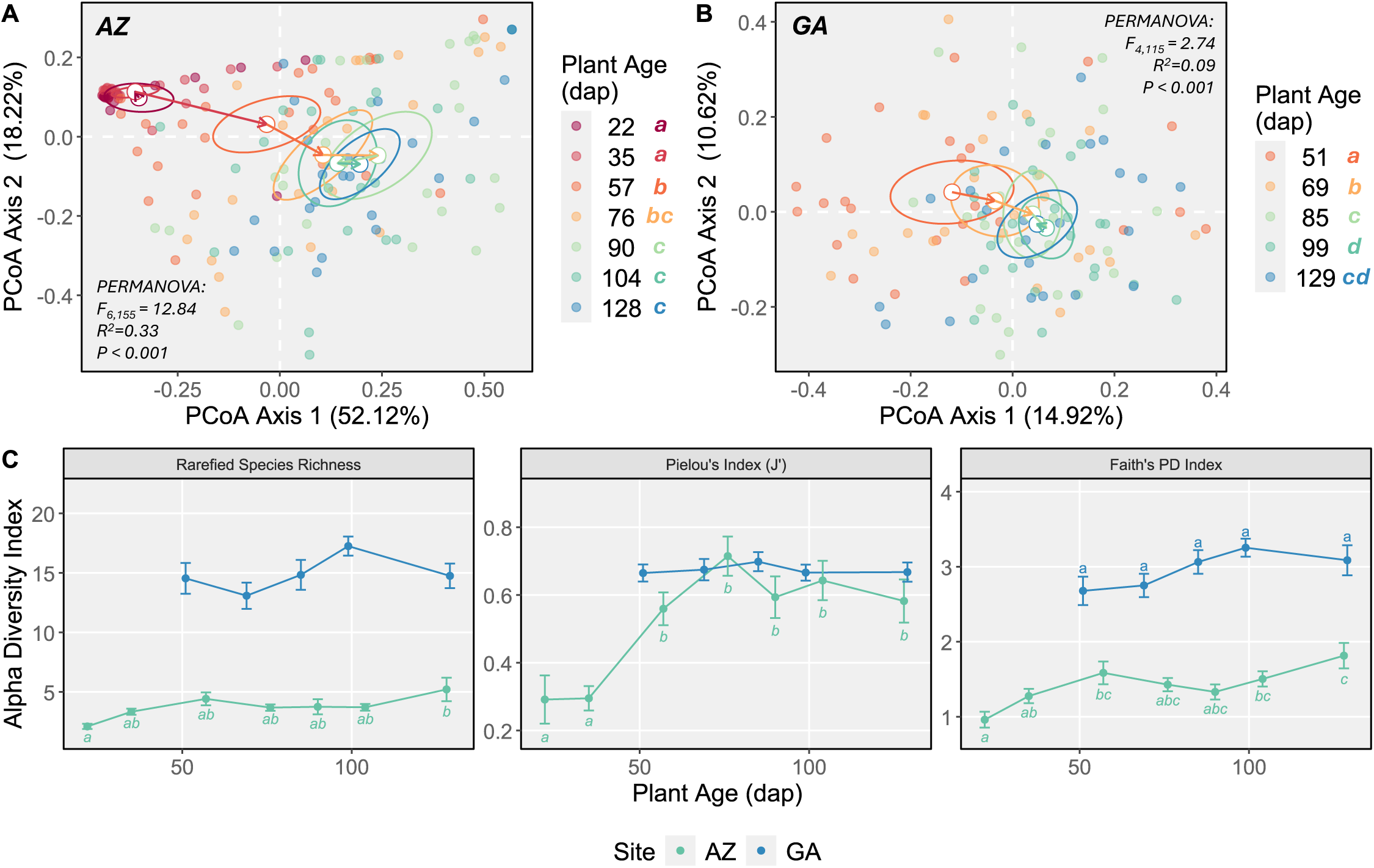
Principal co-ordinate analysis (PCoA) ordinations generated using Bray-Curtis distances between samples for the **A)** AZ and **B)** GA field sites, separated by plant age. Per-group ellipses represent standard error around the mean centroid of samples. Letters next to plant age labels represent groupings of statistical similarity (BH-adjusted p < 0.05) after pairwise PERMANOVA testing following global PERMANOVA model results. **C)** Mean ± SE variation of rarefied species richness, Pielou’s index of diversity, and Faith’s PD in AZ and GA. Letters represent groupings of statistical similarity (BH-adjusted p < 0.05) based on pairwise post-hoc testing of within-site comparisons between plant ages following linear mixed effect models. Replication: AZ, 76 dap n = 22, n=24 for all other plant ages; GA, n=24 per plant age.

The CAP analysis was used to subset a minimal dataset of the ‘most important’ taxa in the magnitude of their contribution to between-genotype community turnover with which we could perform explicit differential abundance tests. Visually, the genotype groups primarily separate along axis 1 of the CAP ordination in both sites, while in GA there is clear genotype separation both within, and between groupings, along axis 2 (**Figure 3 A/B**). *E. etunicata* and *F. mosseae* display the strongest correlation with axis 1 for the AZ CAP ordination and were also found to be significantly different between the three groups (p < 0.05). *E. etunicata* was most abundant in ‘group b’, whereas *F. mosseae* was most abundant in ‘group a’. *E. etunicata* displays the expected intermediate abundances in the ‘mixed group’ subset of genotypes, though *F. mossea* has a lower relative abundance in both the ‘mixed group’ and ‘group b’ relative to ‘group a’ (Supplementary Figure S6). While *D. aurantia* and *De. omaniana* both display a trend of lower mean relative abundances in ‘group b’, and *P. occidentale* has a slightly higher mean relative abundance in ‘group b’ which may contribute to the differentiation between the two groups, these comparisons are not statistically significant (Supplementary Figure S6). For the GA CAP ordination, *Racocetra verrucosa* was the most correlated with axis 1 and was found to be more abundant in ‘group c’ and the ‘mixed group’ than ‘group b’ (Supplementary Figure S7). *P. occidentale* and *P. occultum* were both more abundant in ‘group b’ than in ‘group c’. As was observed for AZ, the remaining independently tested species display smaller variation in abundances between the genotype groups, and while not statistically significant, likely contribute in smaller amounts to the observed turnover between the genotype groups (Supplementary Figure S7).

Genotype differences in aboveground phenotypes of the sorghum genotypes are presented in detail in Supplementary Material Results S1. As was expected, genotypes had contrasting height and biomass phenotypes (Supplementary Figure S10, S11). Genotype-mean phenotype values were positively correlated between sites (Biomass R^2^=0.749, Height R^2^=0.607), but this was only significant (p < 0.05) for biomass, and not for height given sample size limitations indicating a greater potential for heritability in these traits than for the microbiome.

### Temporal dynamics of AMF communities within sites

In AZ, the AMF communities were distinct in the earlier 22 and 35 dap collections with little dispersion between samples, as nearly all roots are heavily colonized by *F. mosseae* (Figure 4a, Figure 5 A/B). Between 57 and 76 dap, the community transitions from this towards a new stable state from 90 dap onwards where we observe increased colonization by later-arriving species including *E. etunicata* and *D. aurantia* as well as other less abundant species (**Figure 4A, Figure 5**). The ordination displays a triangular distribution reflecting the compositional gradients between the three dominant species in AZ. At the later timepoints samples are dispersed between the two alternate tips of the distribution representing *E. etunicata* and *D. aurantia*. The emergent community stability in this case therefore does not come from convergence to a single steady state but is rather reflects a mixture of stochastic colonization by one of the competing and genotype-specific recruitment as previously discussed (Figure 5A, Supplementary Figure S5). Communities in AZ also become moderately more taxonomically and phylogenetically diverse over time. (**Figure 5 C**). Notably, this never results in richness values approaching those seen in GA, though the distribution of taxa within single roots quickly becomes more even to the same extent as we see in GA (**Figure 1 B, Figure 4C**).

**Figure 5.**
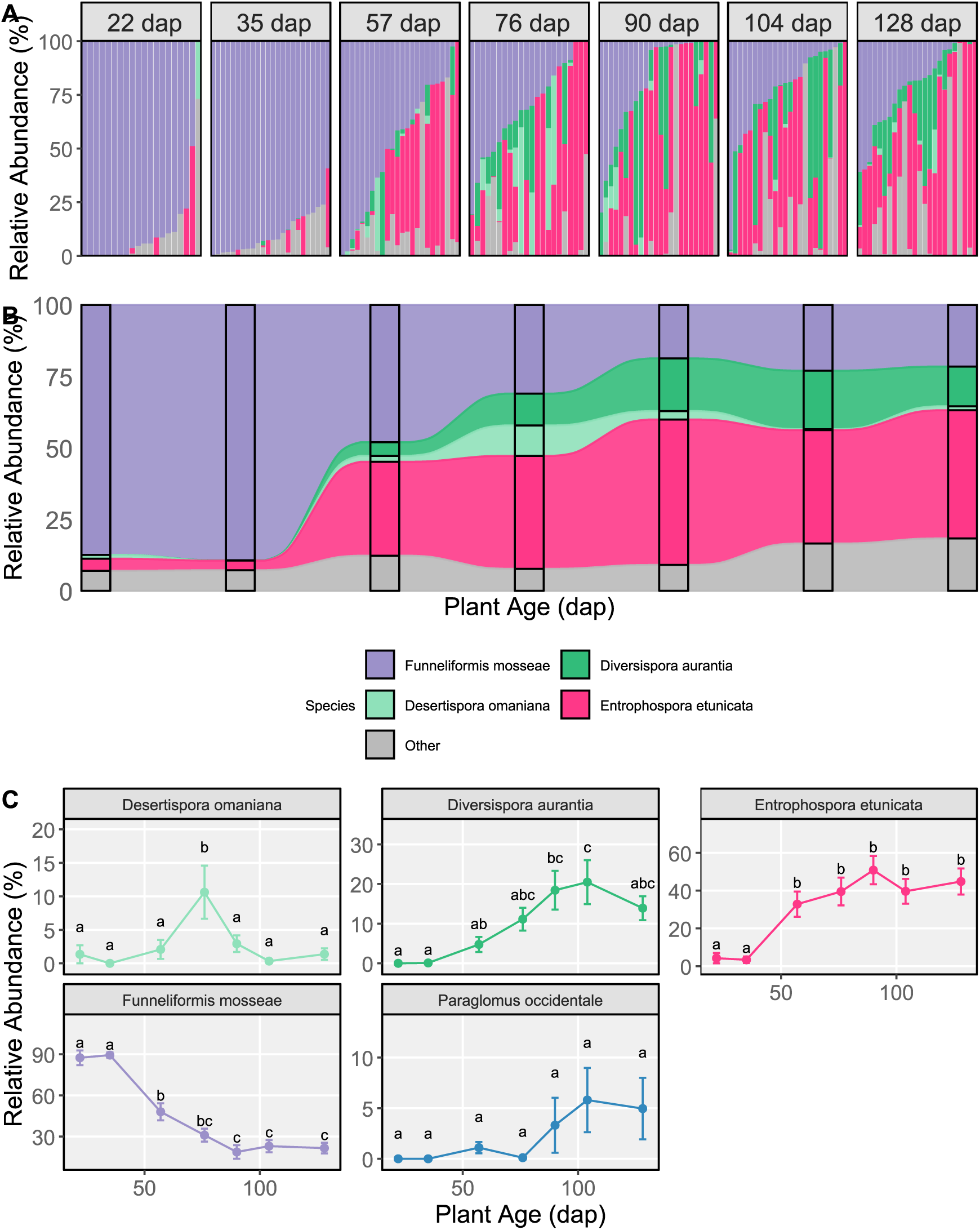
Taxa composition plots of the relative abundance of the four most abundant AMF species in AZ, shown per sample in **A)**, and grouped by plant age in **B)** as mean values. The remaining species are aggregated into one value labeled ‘Other’. **C)** Mean relative abundance ± SE of species found to be significantly different across plant ages through both GAMM model testing. Letters represent groupings of statistical similarity (BH-adjusted p < 0.05) after post-hoc pairwise comparisons. Replication: n = 24 per timepoint.

*Desertispora omaniana* peaks in relative abundance at 76 dap (**Figure 5 C**). This is the same collection point where *D. aurantia* also begins to increase in abundance. Together these two taxa contribute to an increase in the relative abundance of the *Diversisporaceae* family (Supplementary Figure S7), though *D. aurantia* becomes the major representative of this clade. From the GAMM model it was found that *P. occidentale* increases in relative abundance over time, but this was not apparent from the pairwise tests, likely due to high variability (Figure 5 C). As each family is primarily represented by one major species in AZ, family level associations track the patterns observed for these species already described (Supplementary Figure S7).

We did not assess AMF communities in GA at plant ages comparable to the 22 dap and 35 dap collections in AZ, so we cannot ascertain early colonization dynamics for this site. However, we observed a similar pattern as in AZ when considering the similar plant ages assessed, wherein communities showed reduced turnover after 85-99 dap (**Figure 4 B**). Taxonomic diversity also remains relatively consistent over time, though phylogenetic diversity increases slightly (**Figure 4 B**).

*Viscospora viscosa* and *Paraglomus occidentale* remain the most abundant taxa overall across all recorded timepoints (**Figure 6 A, B**). Two species significantly decrease in relative abundance over time (p<0.05, (*Gigaspora margarita, P. occultum*). *F. mosseae* peaks around 69-85 dap before decreasing back to previous abundances (**Figure 6 C**). *Nanoglomus* MT001 increases over time (**Figure 6 C**). At the family level, *Glomeraceae* also increases over time, which cannot be entirely accounted for by the increase in *Nanoglomus* MT001, highlighting that multiple species contribute to the development of the community over time (Supplementary Figure S8). Both the *Paraglomeraceae* and *Gigasporaceae* clades decrease in relative abundance over time, though the *Paraglomeraceae* clade appears to make a small bounce-back at the final observed timepoint (Supplementary Figure S9).

**Figure 6.**
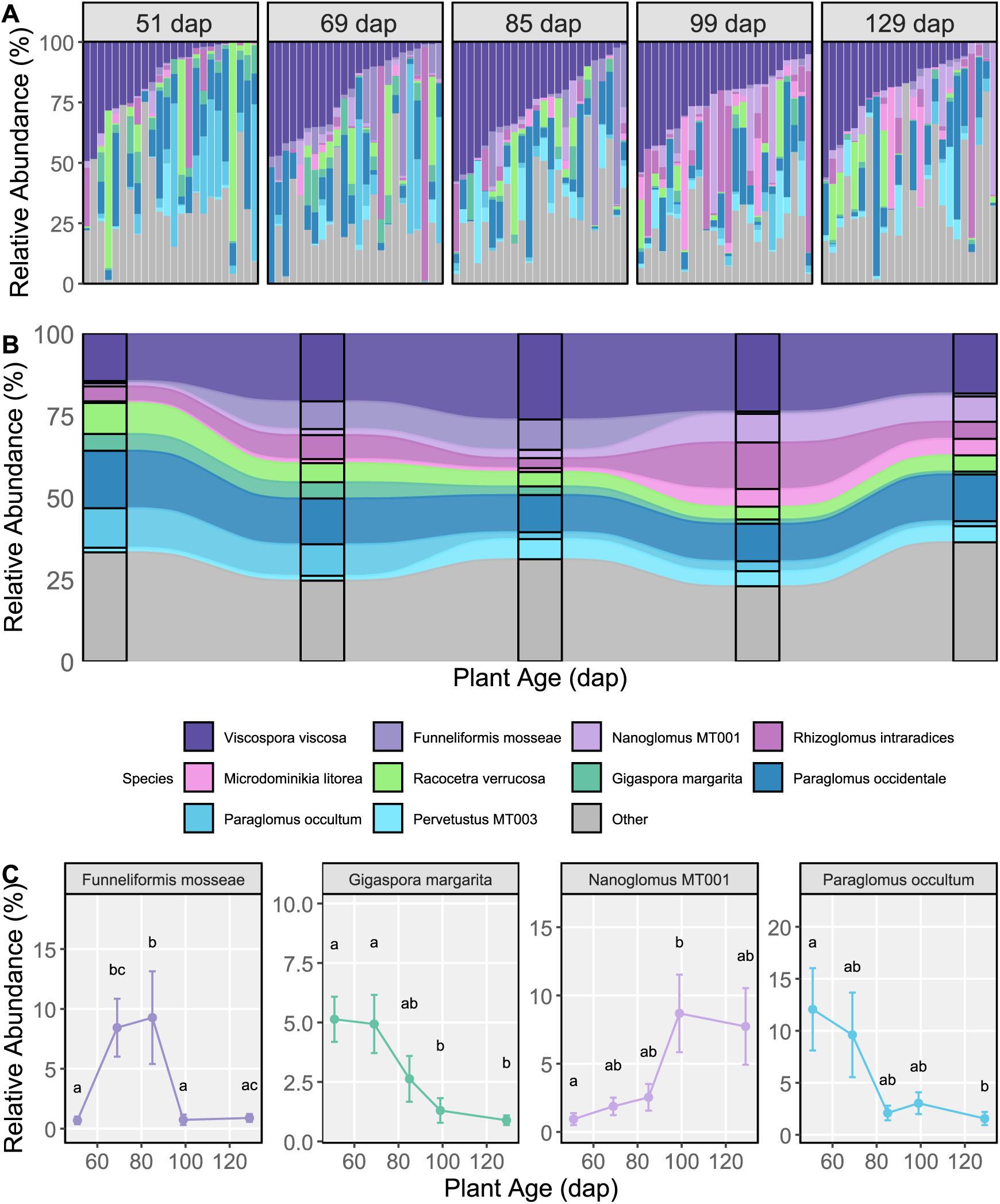
Taxa composition plots of the relative abundance of the ten most abundant AMF species in GA, shown per sample in **A)**, and grouped by plant age in **B)** as mean values. The remaining species are aggregated into one value labeled ‘Other’. **C)** Mean relative abundance ± SE of species found to be significantly different across plant ages through both GAMM model testing. Letters represent groupings of statistical similarity (BH-adjusted p < 0.05) after post-hoc pairwise comparisons. Replication: n = 24 per timepoint.

In further exploring the phylogenetic diversity of samples and the mechanisms underlying community assembly, we examined the standardized effect sizes (SES) of both mean pairwise distances (MPD) between taxa within each sample, and the nearest taxon index (NTI). NTI is a measure of the distance between each taxon and its next closest taxa on the phylogeny. This was performed at the ASV level to capture fine-scale variations, and together these can be informative as to how communities cluster at both a deep (MPD) and shallow (NTI) phylogenetic depth. In AZ, MPD SES values were extremely low during early colonization, reflecting minimal phylogenetic breadth because nearly all ASVs belonged to the dominant *F. mosseae* (**Figure 7 A**). From 57 dap onwards MPD SES exhibits a stark increase and continued to gradually increase until 128 dap where it approached neutrality. This reflects the establishment of *E. etunicata* and *Di. aurantia* in the roots, and gradual increase in phylogenetic breadth within samples as more rare species established within each sample consistent with increasing Faith’s PD (**Figure 7 A, Figure 4 C**). MPD SES in GA remained moderately negative across all observations, reflecting consistent clustering at deeper phylogenetic levels and limited changes in phylogenetic breadth despite consistent turnover in composition, though we notably do not have analogous collections in GA to those in AZ that were dominated by *F. mosseae* (Figure 8A). The negative MPD SES reflects high abundances of taxa belonging to either the *Glomeraceae* or *Paraglomeraceae* family clades. At both sites NTI SES was consistently slightly negative (Figure 8B), denoting weak ‘environmental filtering’ favoring tip-level clustering of closely related taxa. When taken in the context of our between-genotype comparisons, these patterns are consistent with plant genotype effects acting as the ‘environmental’ filter favoring compatible AMF clades.

**Figure 7.**
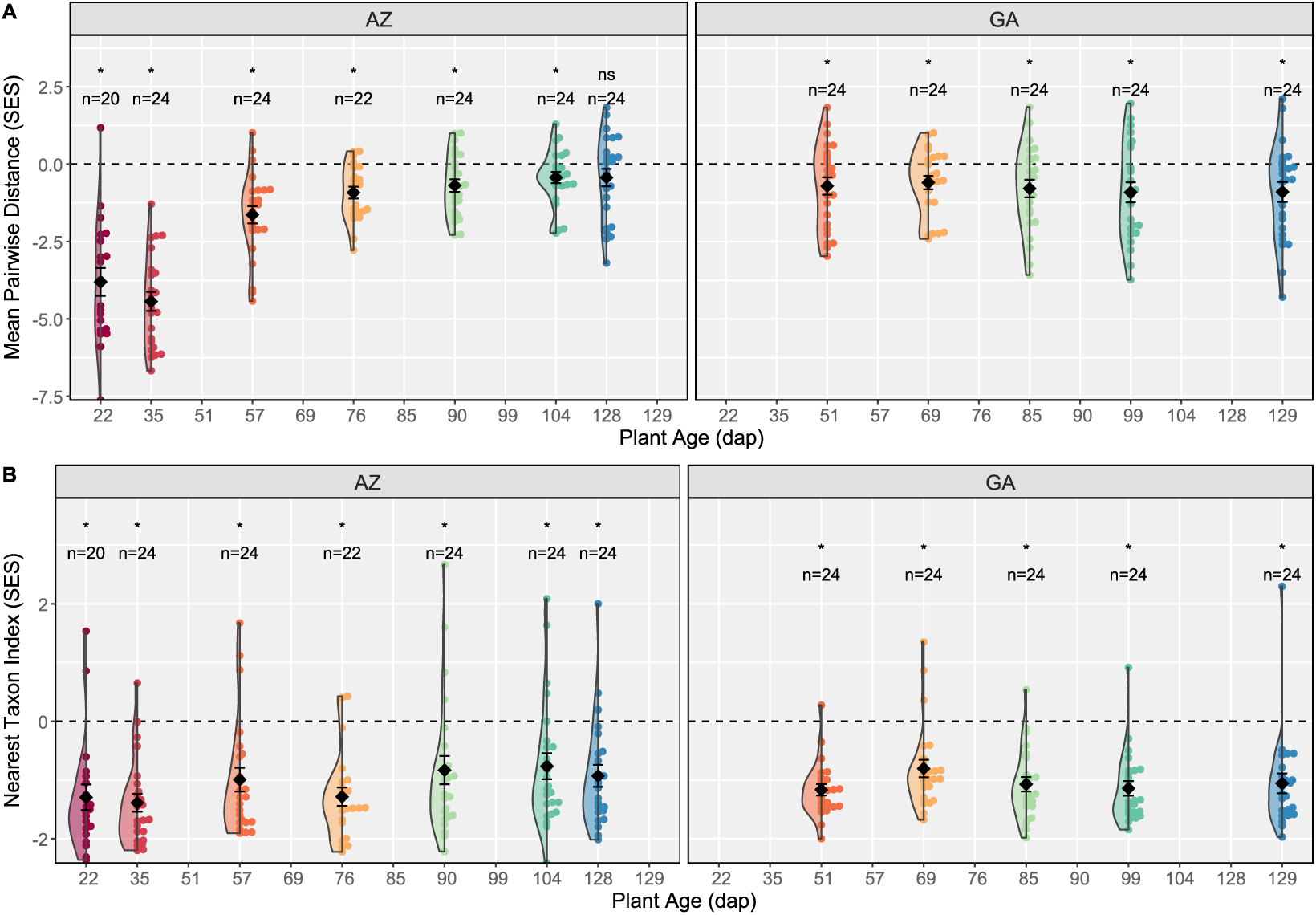
A. Combined violin / bee swarm plots displaying the variation of Mean Pairwise Distances (MDP) standardized effect sizes (SES) in **A)** AZ and **B)** GA, and Nearest Taxon Index (NTI) SES in **C** AZ and **D)** GA. Values are separated by plant age. Mean ± SE are also displayed for all values. Deviation from zero was tested by t-test individually for each plant age per site, and asterisks denote where this was significant (p < 0.05).

## Discussion

Our findings are consistent with expected AMF community patterns related to precipitation / temperature gradients (House & Bever, 2018; Sousa et al., 2022), and soil edaphic properties (Oehl et al., 2017; Verbruggen et al., 2012) affecting both community diversity and assembly. We observed lower species richness, evenness and phylogenetic diversity in AZ relative to GA, reflecting the highly distinct environmental contexts of the two sites. Soil P and Ca were 58-times and 9.7-times higher respectively in AZ than GA. Such high nutrient conditions are regularly associated with low-species richness and evenness due to through strong selection pressure for taxa that handle extreme conditions and rapidly utilize the nutrient source (Alguacil et al., 2010; Ma et al., 2020; Revillini et al., 2019). AZ is dominated by three taxa with > 10% dataset-wide relative abundance-*F. mosseae* (28.4%)*, E. etunicata* (41.6%) and *D. aurantia* (13.79) with a combined occupancy of 67.8%. Only seven species are present in > 1% abundances. In GA, *. viscosa* and *P. occidentale* are present > 10% with a combined occupancy of 34.4% and 17 additional species are present in > 1% abundances. There were a further 62 rare species (< 1%) present, 20 of which are completely absent in AZ, which has no unique species present.

The low richness and phylogenetic diversity in AZ points to strong environmental selection pressure for ruderal and stress tolerant characteristics, which are displayed by the subset of taxa present in high abundances in AZ (Chagnon et al., 2013; Millar & Bennett, 2016). Members of *Glomeraceae* (historically incorporating *Claroideoglomeraceae syn. Entrophosporaceae*) are typically considered ruderal due to their fast hyphal turnover and interconnectedness (Voets et al., 2006) and a preferential allocation of biomass to root colonization (Alguacil et al., 2010; Weber et al., 2019). *F. mosseae* and *E. etunicata* belong to *Glomeraceae* and *Entrophosporaceae* respectively, and have cosmopolitan distributions (Stürmer et al., 2018), though they are notably in low abundance in GA relative to AZ. The ruderal lifestyle of both species is conserved across a variety of habitats, through moderate to high root-colonization and spore abundances in habitats including high-disturbance agricultural soils (Li et al., 2007; Oehl et al., 2017), hypersaline soils (Silvani et al., 2017), and drought-prone arid environments (Mohammad et al., 2003) such as those in AZ. *Diversispora* species are frequently associated with stress-tolerant characteristics through the preferential production of persistent soil hyphae (Chagnon et al., 2013; Weber et al., 2019) and resistance to abiotic stress. *Diversispora* is prevalent in arid desert environments, and those with strong wet/dry seasons (Bahadur et al., 2019; Raya Montaño et al., 2019; Symanczik et al., 2014, 2015; Vasar et al., 2021), as well as early successional communities that are analogous to the AZ field site (Sikes et al., 2012).

Along with nutrient availability, plant host presence presents an additional selection pressure on AMF taxa in the AZ site, towards species capable of forming long-lasting resilient spores that can persist in the absence of a regular host. Sorghum was cultivated in multi-year bare soil plots in AZ, contrasting the freshly cleared old-field site of GA. We posit that a native AMF inoculation source would be present across robust networks of existing plant roots, hyphae, and spores in GA, whereas in AZ the resident AMF would rely primarily on the spore bank for persistence between crop cycles which would further bias communities towards *Glomeraceae* and *Entrophosporaceae* due to their capacity to form large spore volumes (Hempel et al., 2007).

Sorghum cultivated in AZ exhibited a markedly higher biomass at the final harvest relative to that in GA, while having lower AMF species richness, evenness and phylogenetic diversity. This is not surprising given that high P content decouples the microbe-mediated nutrient acquisition via AMF from plant nutrition (Hodge et al., 2010). Sorghum is also well adapted to arid areas (Tari et al., 2013) and, in the water depauperate AZ system, is largely released from both abiotic and biotic pressures on yield otherwise present in GA, including weed competition (Zimdahl, 2008), pathogen loads and pests (Tscharntke et al., 2005). The notion that AMF biodiversity intrinsically begets nutritional benefit to plants has also been shown to be an oversimplification that does not hold up well under many experimental conditions (Powell & Rillig, 2018). In a study comparing monoculture vs polyculture, (Antoninka et al., 2011) showed that polyculture resulted in a reduction of AMF diversity and soil-exploring extra-radical hyphal density, but this farming strategy was associated with an increase in both root and shoot biomass, possibly driven by host plant selection for highly compatible AMF. Drawing specific associations between microbiome assembly and plant yields would require further within-site experimental manipulations to explicitly test the relative importance of species richness, diversity and composition on plant yields.

In contrast to AZ where low phylogenetic diversity selects for groups with a potentially narrow range of life history traits (Maherali & Klironomos, 2007; Powell et al., 2009), GA has varied representation of multiple species from clades associated with both root and soil colonization and which may display contrasting niche separation within this environment. While AZ is dominated by a single *Funneliformis* species, the most indicative genera of GA from this clade are *Nanoglomus*, *Dominikia, Microdominikia*, *iscospora*, and *Rhi oglomus*. As a very closely related genus-*iscospora* may occupy a similar niche to *Funneliformis* along with *Rhi oglomus*, though the other groups are relatively unexplored. GA is also low in *Diversisporacea*, instead containing taxa from the closely related *Gigasporaceae* clade (*Racocetra*, *Cetraspora*, *Gigaspora*, *Scutellaspora*) and *Paraglomeraceae*. Like Diversispora, *Paraglomeraceae* and *Gigasporaceae* are both preferentially occupy the soil niche through hyphal development (Hempel et al., 2007) that may be selected for by the plant host in low nutrient conditions like GA (Kiers et al., 2011; Verbruggen & Toby Kiers, 2010). Gigasporaceae also invests highly in soil hyphae, though this tends to be short transport hyphae (Hart and Reader 2005, Souza et al 2005) with low connectivity that may indicate a lack of stress tolerance.

At both sites we see evidence of host filtering that is evident from our between-genotype assessments of root community composition, species abundances between genotype-groups, and our assessment of phylogenetic patterns across time. Consistently negative NTI values were observed at both sites, and can be associated with competitive exclusion and ‘environmental’ filtering (in this case, the host being an environment). Host-mediated microbiomes were less detectable in AZ than GA despite selecting from a genetically diverse population of sorghum genotypes. The source community available in AZ likely provides less opportunity for host-selections, evidenced by both reduced visual discrimination between communities of each genotype and a greater number of genotypes belonging to the ‘mixed’ group (i.e., not recruiting a particularly unique community) in AZ than GA. This initial finding on a limited subset is somewhat obscured by the signal-to-noise ratio associated with sampling a small number of contrasting genotypes. We suggest that if we were to expand our assessment to the whole BAP panel we could better resolve broad patterns of host genetics and explore the heritability of AMF community assembly through techniques such as microbiome GWAS analysis, as has been done for bacteria in other systems (Deng et al., 2021; Edwards et al., 2023; Sutherland et al., 2022). We do, however, highlight that the degree of top-down genotypic control of community assembly may itself be a highly plastic trait, and highly dependent on the source environment and community selected for.

Our results contrast with those of (Frew, Heuck, and Aguilar-Trigueros 2023) who observed that *Gigasporaceae* and *Diversisporaceae* displayed opposing preferences to phosphorus availability to what we observed. Finding consistent ecological patterns across different sites and contexts is complicated by the plethora of other contributing factors often not measured, but this does demonstrate that phosphorus status is not a sole factor determining the abundance of these two clades. (Frew et al., 2023) also observed evidence of phylogenetic overdispersion under low P, and strong environmental filtering under high P, which they determined as evidence of host-plant filtering based on the assessment of nearest taxon index and mean pairwise distances between taxa. In contrast we found low evidence of overdispersion, instead finding environmental structuring being the prevailing structuring force at both sites. As similar patterns of phylogenetic dispersion can have multiple underlying mechanisms, we cautiously propose that this is additional evidence of host-filtering rather than through a different filter (e.g., edaphic properties) in the context of our study, as we observe additional evidence of contrasting community phenotypes across our genetically diverse sorghum hosts.

A previous study (Gao et al., 2019) observed strong patterns of AMF temporal succession when assessed at the OTU level from a field in southern California USA also considering *Sorghum*. While the resolution of analysis differs, we found similar patterns at both sites in our own study, though they are best exemplified by the AZ site where we had observations of earlier colonization. In their field study (Gao et al., 2019) found that early dominant OTUs of *Rhi ophagus* (syn. *Rhi oglomus*) and *Claroideoglomus* (syn. *Entrophospora*) were rapidly displaced by initially rare OTUs also belonging to high root colonizers including *Rhi oglomus* taxa, *Glomus*, and *Funneliformis*. In our study, it was *F.mosseae* that began as the high abundance taxon, likely driven by priority effects because of its ability to rapidly colonize young roots (Hart & Reader, 2002; Werner & Kiers, 2015). In this case it was rapidly displaced by *E. etunicata*.. In both studies diversity also increased over time, though this was again to a more significant extent in GA. Stronger diversity patterns and intra-species patterns of succession would be more evident if we considered them at the out level as well, but that was not within the remit of this study.

There is an intersection between our temporal and between-genotype assessments wherein in AZ some genotypes continued to be more strongly associated with *F. mosseae* at later timepoints even as the site-wide trend was for this species to be replaced more by other species. PI 92270 and PI 452971 displayed consistently different associations with *F. mosseae* regardless of the site, though in GA this is somewhat driven by a few samples with high colonization by this taxon in PI 452971. The strong priority effects leading to ubiquitous *F. mosseae* presence at these early timepoints may have been countered by some strong competitive advantage of *E. etunicata* at the AZ site, yet *F. mosseae* persisted in part possibly due to its compatibility with some of the *Sorghum* genotypes.

We modified the FLR3-FLR4 amplicon approach previously used in AMF studies, substituting the FLR4 primer sequence for FLR2 to avoid the known amplification biases associated with FLR4 (Gamper et al., 2009; Kohout et al., 2014). While this amplicon enabled us to capture a broad spectrum of AMF across families, it presented additional bioinformatics challenges associated with handling non-specific amplification and required multiple re-sequencing efforts. We therefore would recommend employing a nested PCR approach to better target AMF exclusively or simply utilizing long-read sequencing technologies to capture the full LSU region (and beyond). Full LSU analysis would make better use of recently developed community database and phylogenetic resources (Delavaux et al., 2025; Tedersoo, Hosseyni Moghaddam, et al., 2024).

While there are resolution-based drawbacks to SSU sequencing (Schlaeppi et al., 2016), the use of phylogenetically informed virtual taxa (VTs) circumvents issues arising from an ever-evolving understanding of AMF taxonomic delineations (e.g., (Błaszkowski et al., 2022; Sieverding et al., 2015)) and provides a consistent basis for cross-experimental comparisons. As more sequences are generated such methodology may be better applied to LSU data using foundational resources such as those recently generated by (Delavaux et al., 2024; Tedersoo, Hosseyni Moghaddam, et al., 2024) to better marry morphological ID with molecular taxonomy. Currently, achieving accurate and consistent identification at the species level using LSU taxonomy presents novel challenges. In our study, the taxonomic assignments based on implementing the QIIME2 naïve Bayesian classifying algorithm against the EUKARYOME LSU database mostly show good agreement with our LSU phylogeny (created through the decoration of the Delavaux backbone tree ‘ *16_LSUDB_2024’* (Delavaux et al., 2024)). It does, however, highlight some inconsistencies between historic taxonomic assignments and LSU phylogeny that become apparent from comparing EUKARYOME assignments to the Delavaux backbone sequences, many of which are derived from curated INVAM isolates. As an example, three INVAM isolates assigned to *P. occultum,* when decorated with our environmental ASVs, fall within three separate clades of environmental sequences assigned to *P. occultum* (INVAM CR402)*, P. occidentale* (INVAM VA102A), and *P. laccatum* (INVAM SF123) through EUKARYOME feature classification. This split is not apparent from the backbone phylogeny and becomes emergent only when a greater number of environmental sequences are added. Such discrepancies may either be a limitation of the LSU barcode region through considering intra-specific diversity of sequence clustering (Thiéry et al., 2012), or taxonomy labeling. Given this, we cautiously assign our molecular taxa only to better understand our specific study system and hope that as sequencing efforts continue such assignments become superseded. We also supply all of our input data as ASVs to facilitate any future re-annotations as long-read and whole genome sequencing (Corradi et al., 2024) taxonomy efforts continue.

### Conclusion

In this study, we explored AMF community dynamics as an extended phenotype of the host plant, in the context of environmental selection pressures and intra-species diversity of microbiome assembly for the crop plant *S. bicolor* ((Favela et al., 2023; Johnson & Gibson, 2020)). Through this, we begin to unpack some of the multifaceted determinants of AM fungal community assembly, revealing how environmental filtering, host-plant genotype, and temporal dynamics can shape the composition of root-colonizing AMF. We found that the local environmental context was the major determinant of AMF community composition, acting as a filter for the source community available, as has been previously observed in very different environmental circumstances (Opik et al., 2010; Stürmer et al., 2018) and without our level of AMF species resolution. This context-dependent filtering appears to have downstream effects on genotype level assembly and on the ecological processes shaping AMF community assembly over time. The host GxE interaction was also found to significantly contribute to community composition, though this was minor in the context of overwhelming environmental filtering. We chose our sorghum genotypes to capture a breadth of underlying genetic diversity based on population sub-structure as a coarse examination of the potential spectrum of AMF associations to guide future research. Explicitly leveraging host-genotype to manipulate crop microbiomes is a rapidly developing topic that has seen increased interest in recent years (Chaluvadi & Bennetzen, 2018; Clouse & Wagner, 2021; Oyserman et al., 2020) as an important tool in maximizing microbiome contribution to sustainable agricultural systems. We demonstrate from the GxE outcomes of this study that the innate microbiomes at different sites may be conducive to leveraging host-genetic capital. This provides an important baseline to establishing sites that will most benefit from specific intervention to foster a maximally useful AMF consortia that contribute to sustainable agriculture goals.

## Acknowledgments

DNA Sequencing was performed by the UGA Genomics and Bioinformatics Core (GGBC, UG Athens, GA, RRID:SCR_010994). Soil chemistry was performed by the UGA Extension Agricultural and Environmental Services Laboratories. This research was supported by the US Department of Energy grant DE-SC0021386.

## Author Contributions (CRediT)

**PBC:** Conceptualization, Methodology, Formal Analysis, Investigation, Data Curation, Writing-Original Draft, Project Administration **THP:** Conceptualization, Methodology, Investigation, Writing-Review and Editing, Project Administration **BJL:** Methodology, Investigation, Writing – Review and Editing, Project Administration **AKB:** Investigation, Writing-Review and Editing. **SRM:** Investigation, Writing-Review and Editing. **SS:** Conceptualization, Methodology, Investigation, Writing-Review and Editing. **NCJ:** Conceptualization, Methodology, Writing-Review and Editing, Supervision. **KMD:** Conceptualization, Methodology, Writing-Review and Editing, Supervision. **JLB:** Conceptualization, Methodology, Writing-Review and Editing, Supervision.

## Data Availability

Raw forward and reverse sequences per sample are available under the NCBI Sequence Read Archive (BioProject *TBD*). Denoised and filtered ASV sequences used for experimental analysis are available through NCBI GenBank (Accessions: *TBD*). The complete bioinformatics workflow used to process ASVs and generate phylogenies, along with the R script used for statistical analysis and visualization [will be] available through GitHub (*TBD*).

---

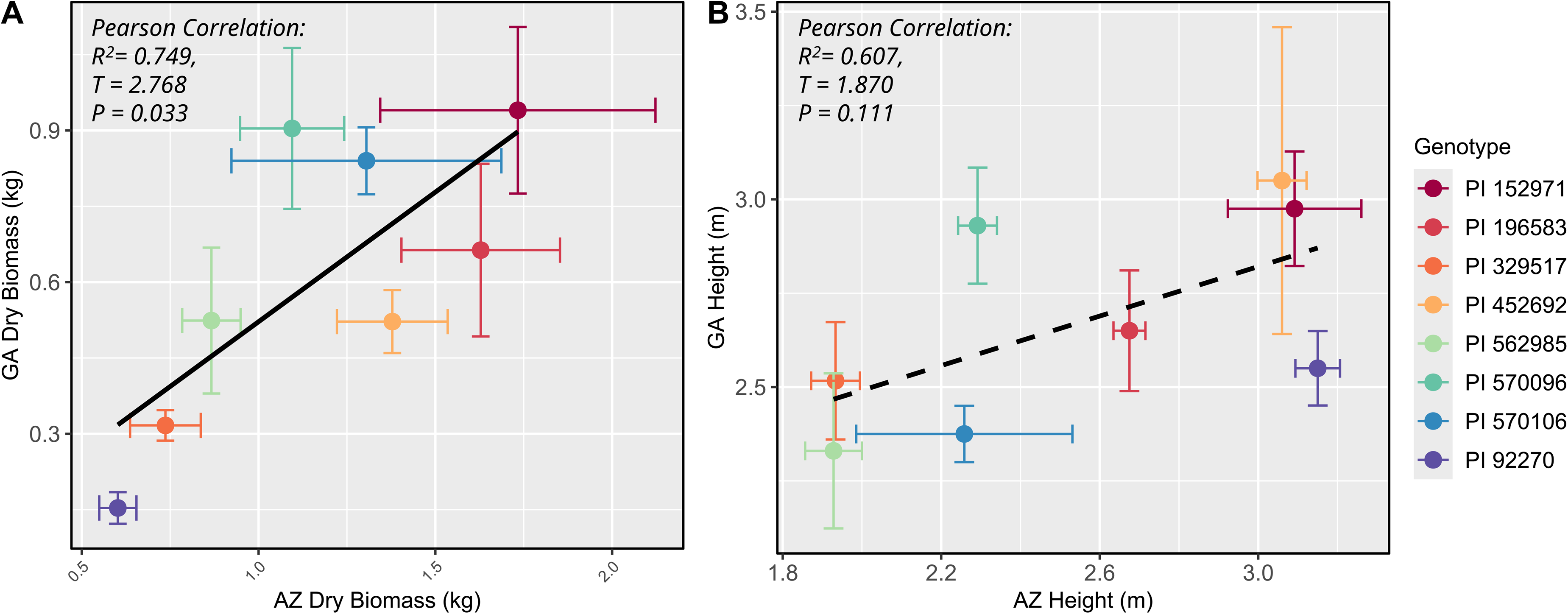

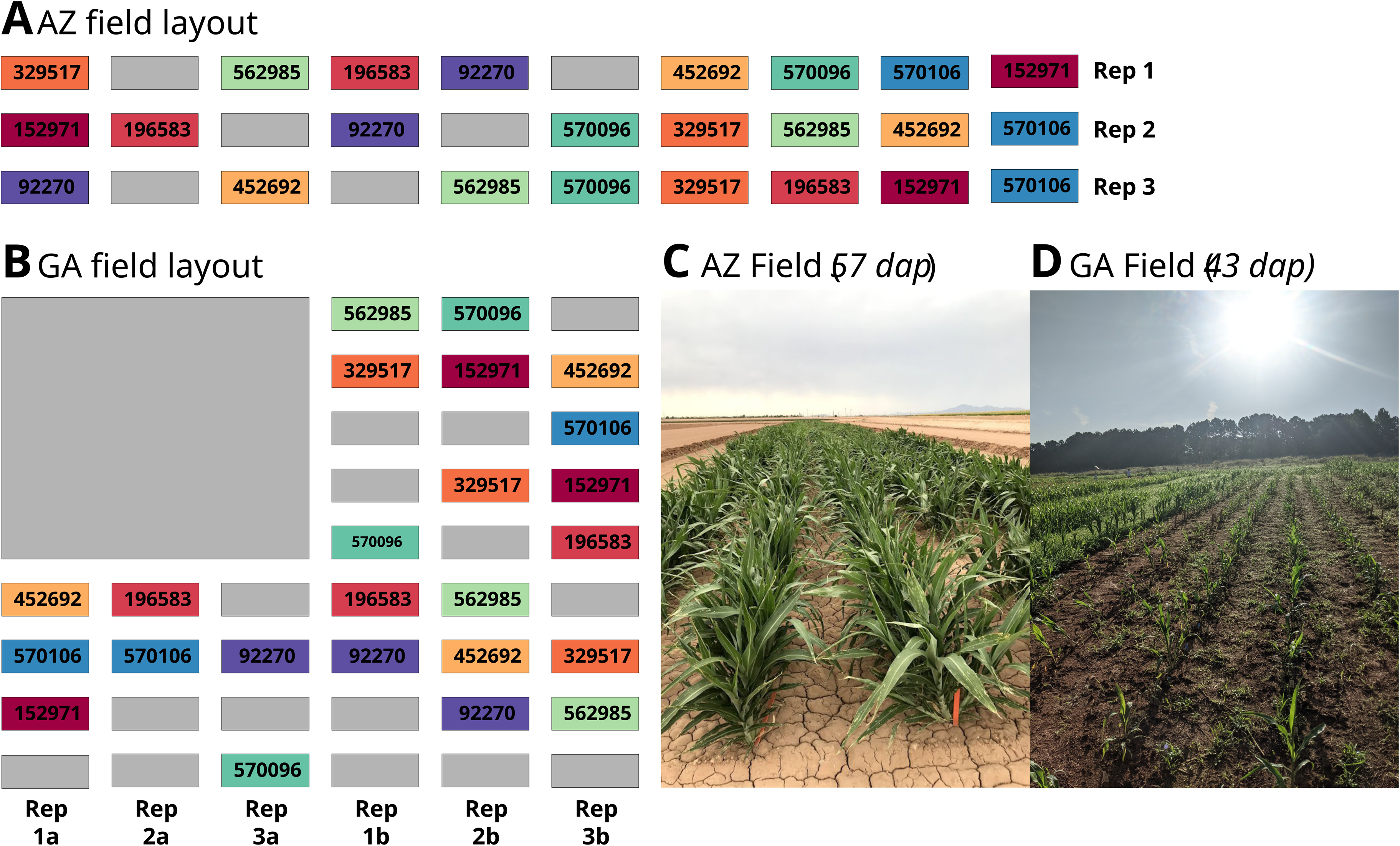

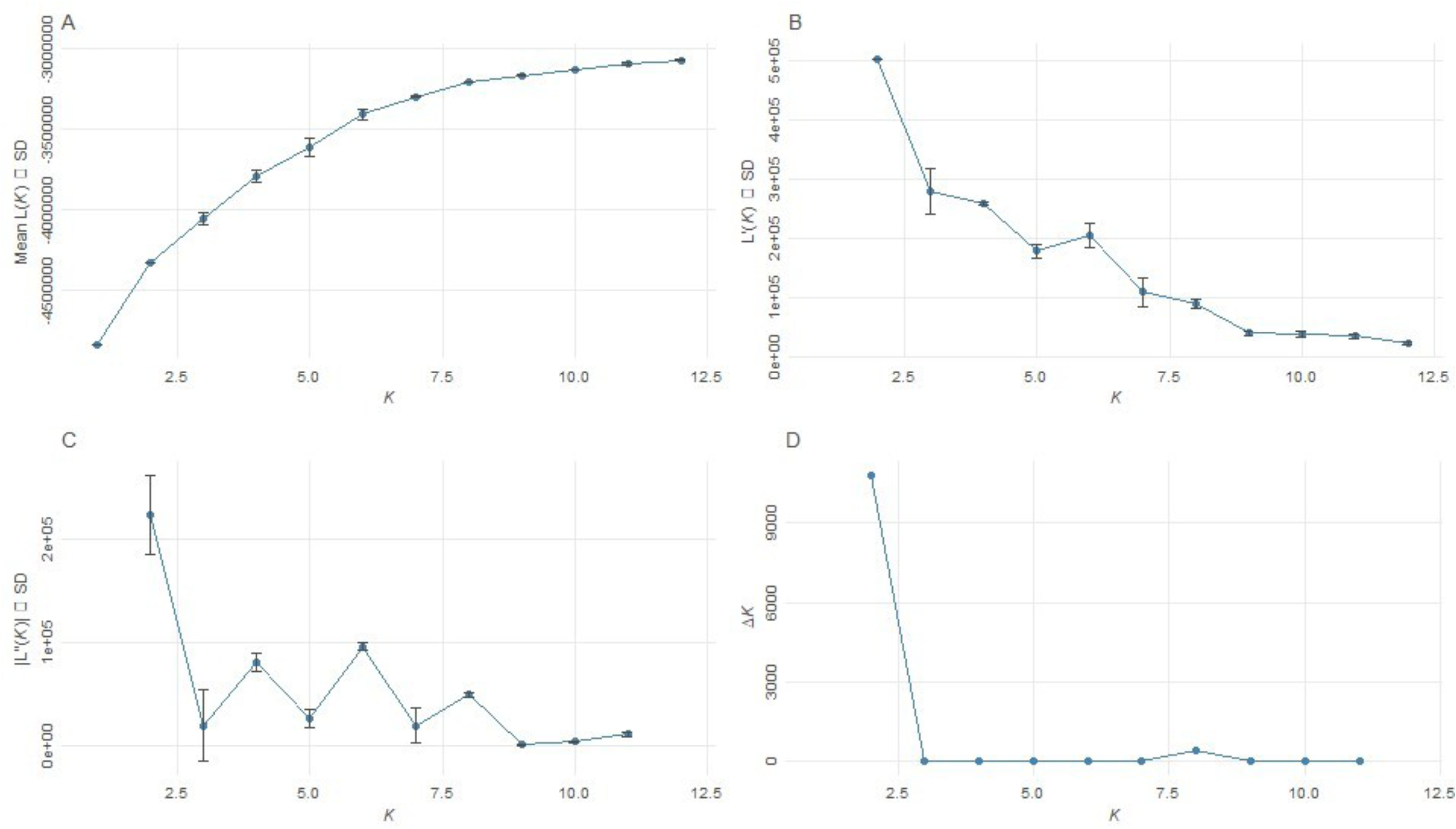

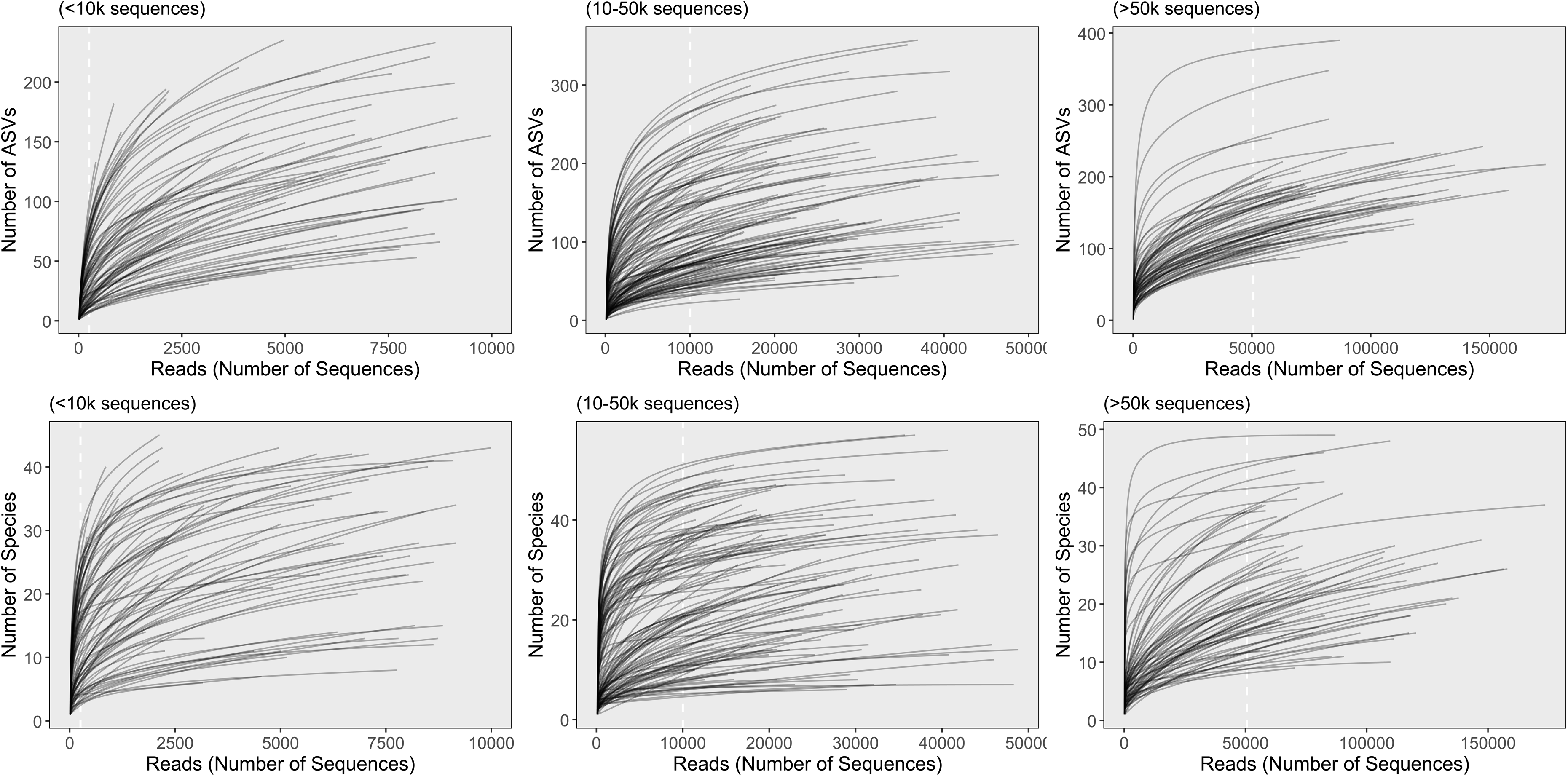

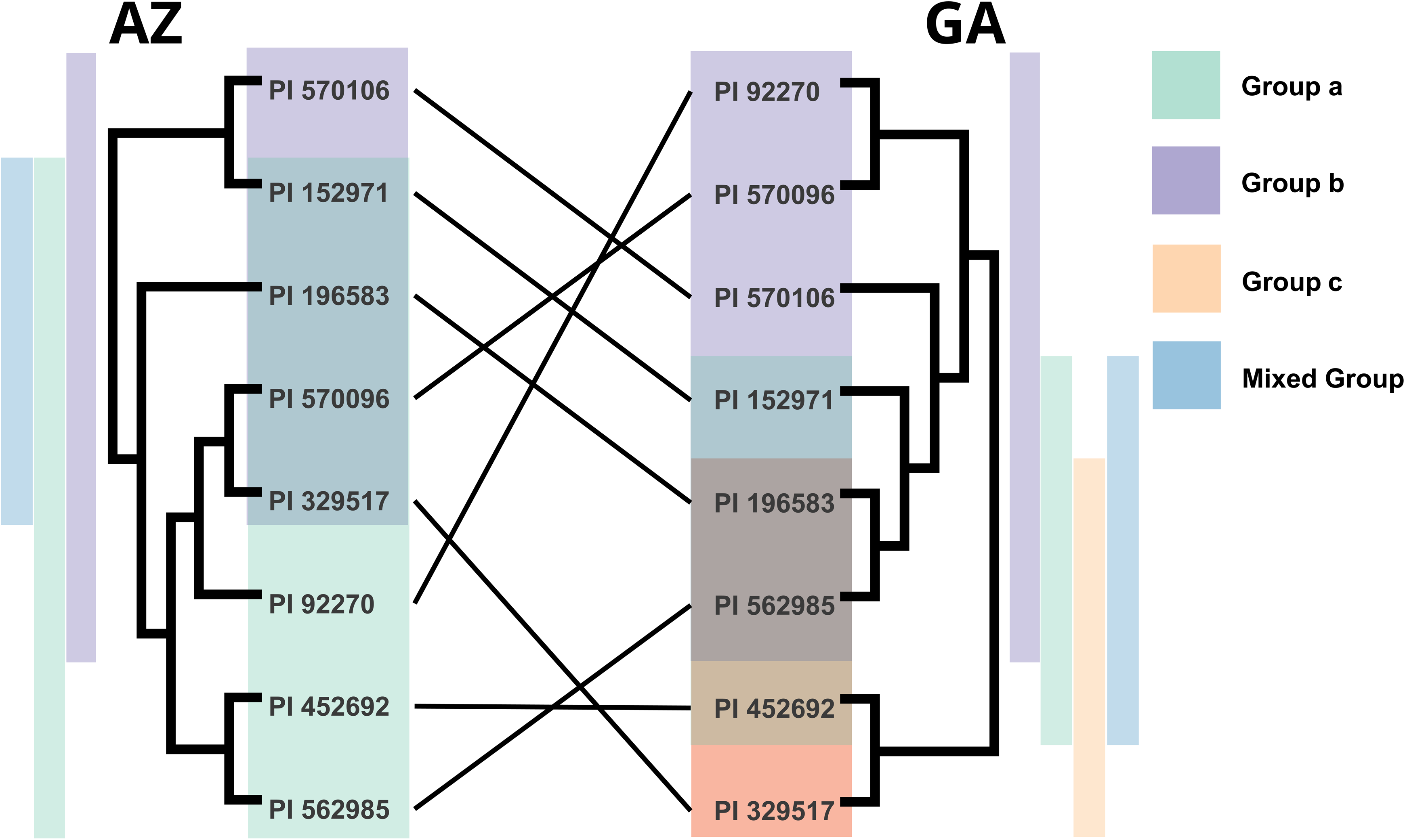

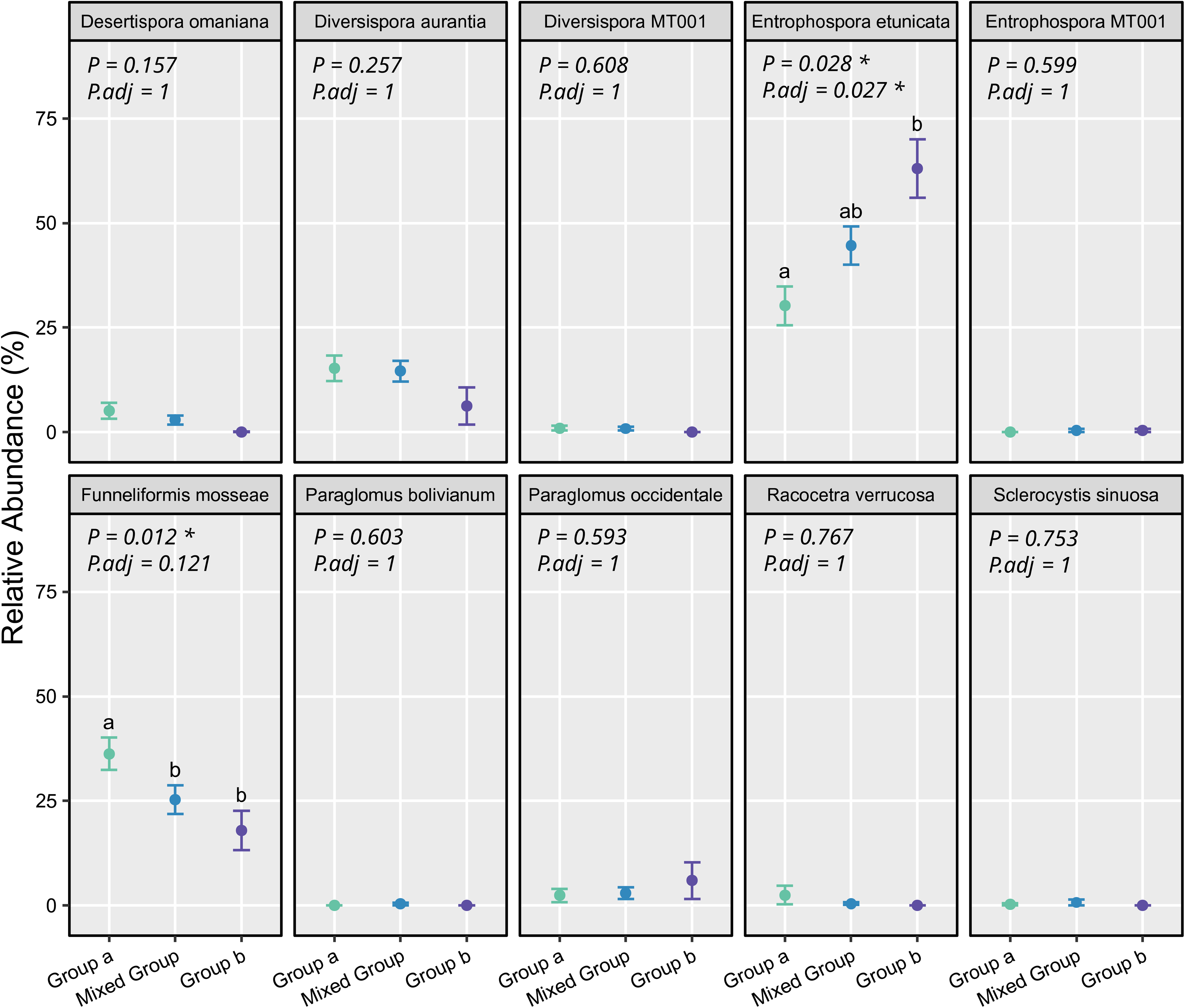

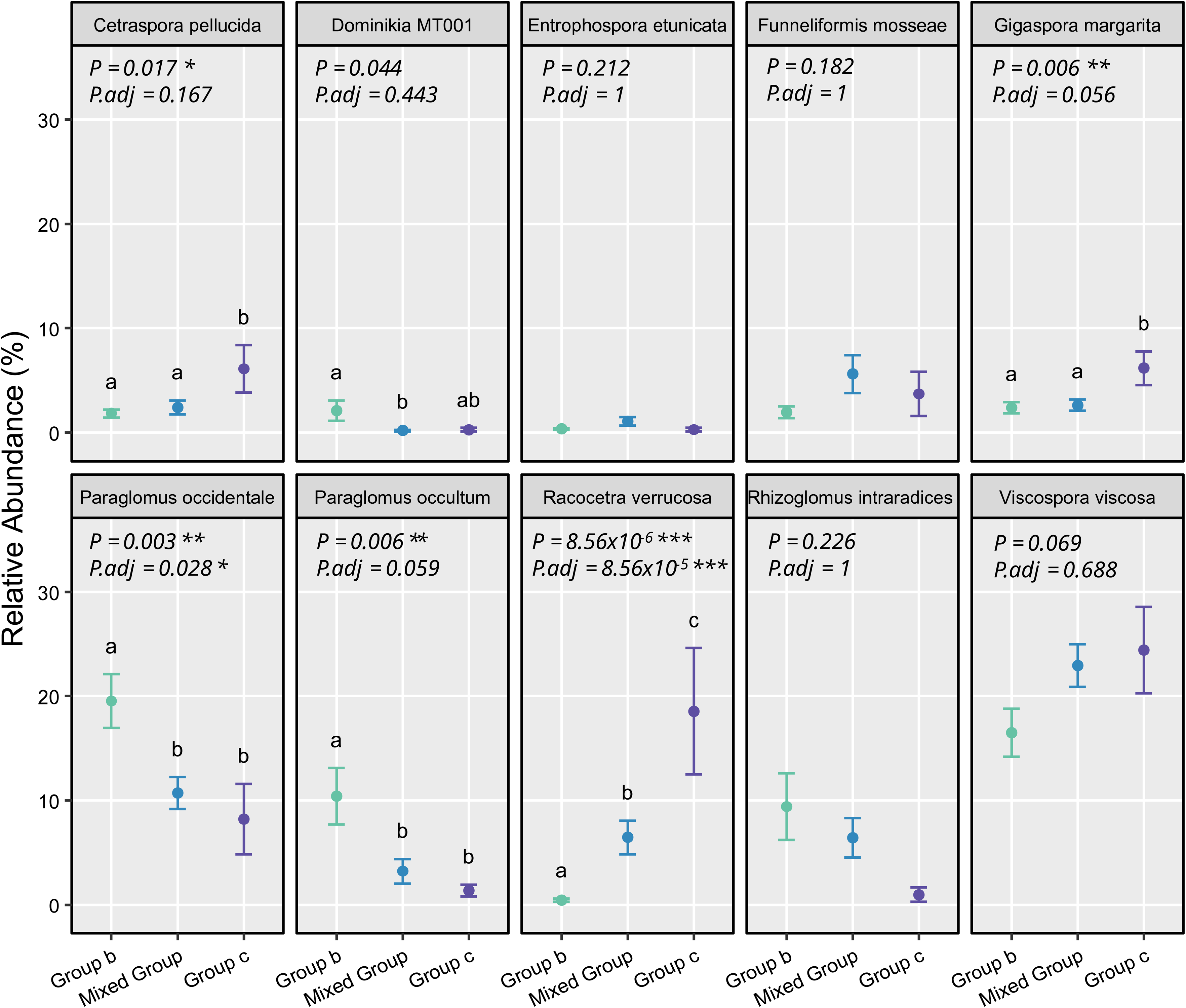

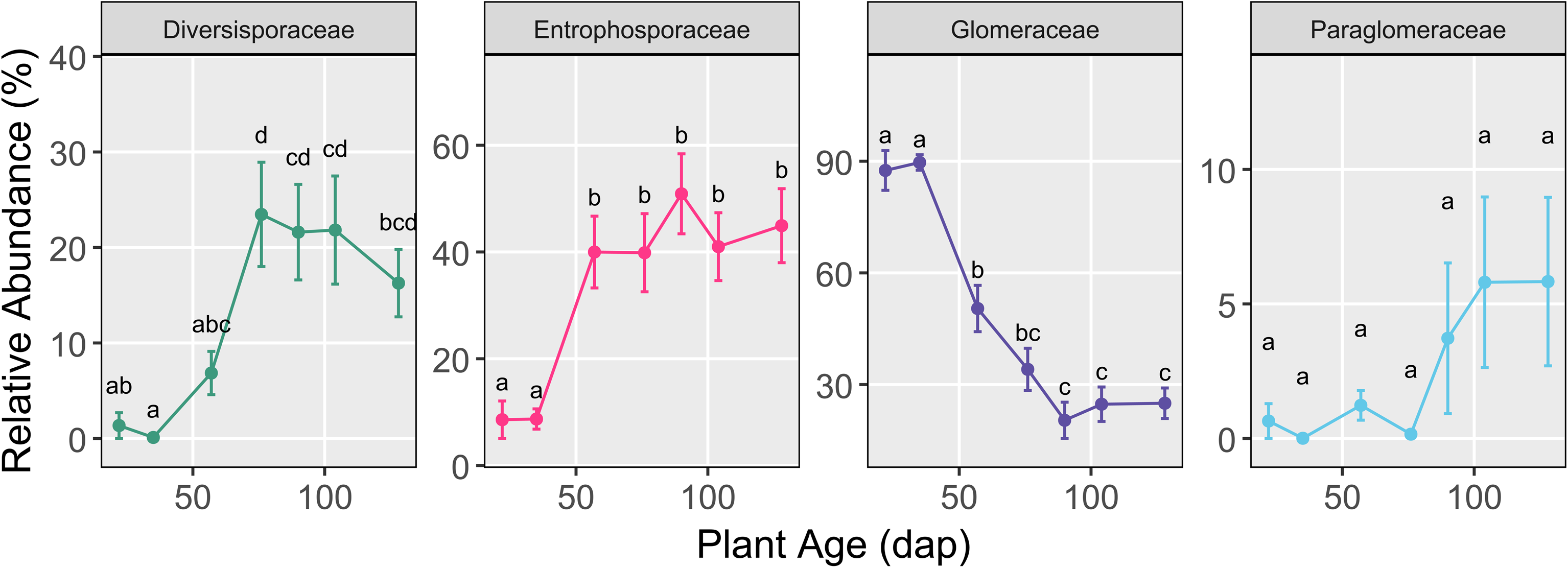

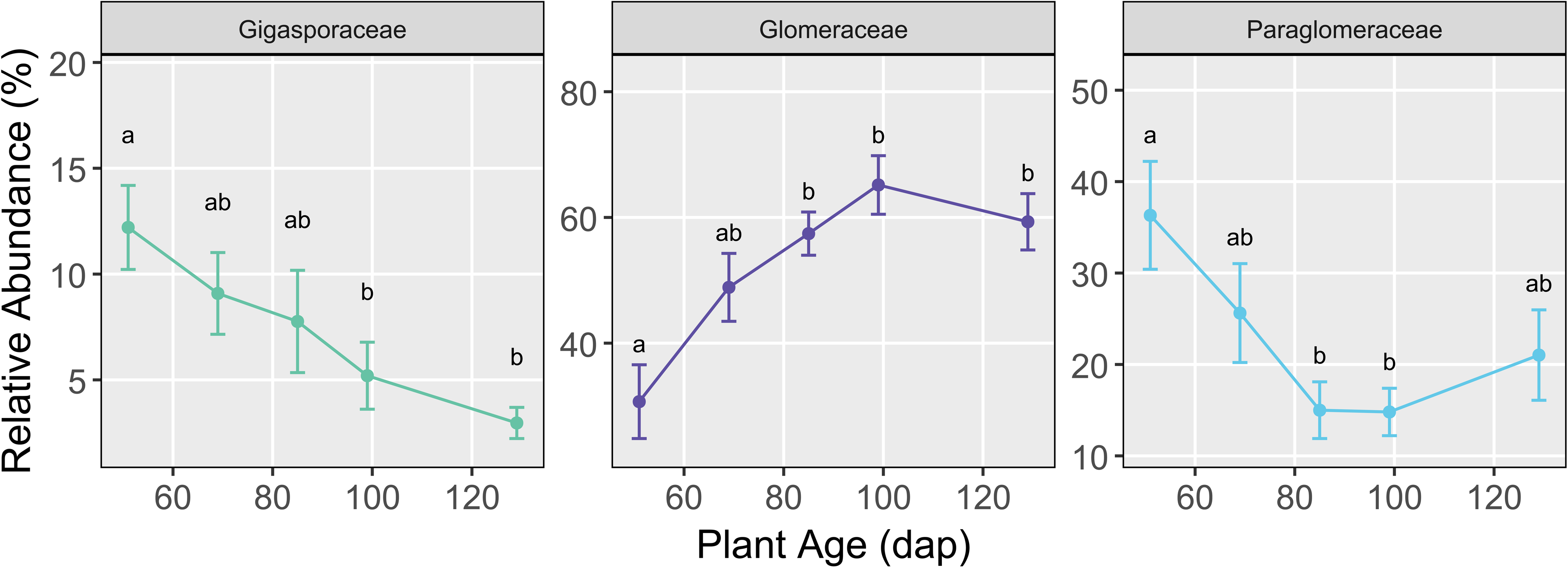

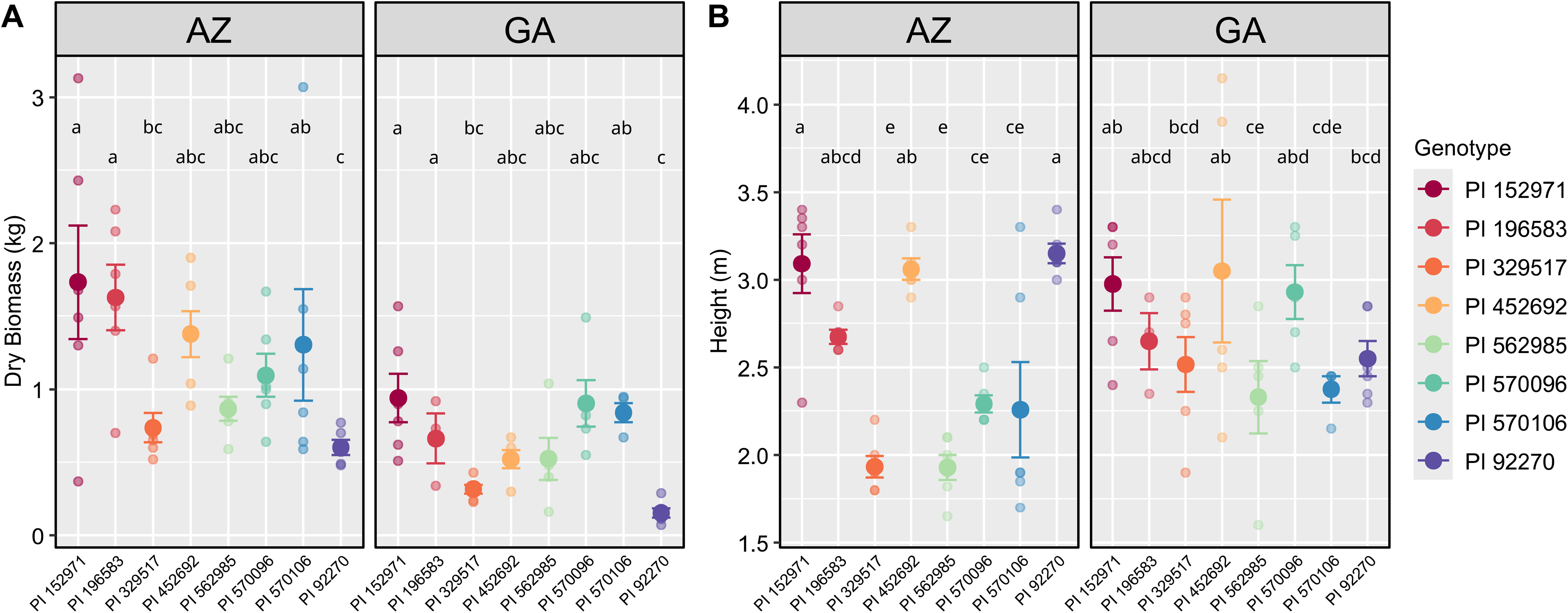

