## Supplementary figures and images for "Exploring eco-evolutionary and temporal patterns of arbuscular mycorrhizal fungal communities colonizing *Sorghum bicolor* across sites of contrasting land use history and climate"

### Supplementary File S2

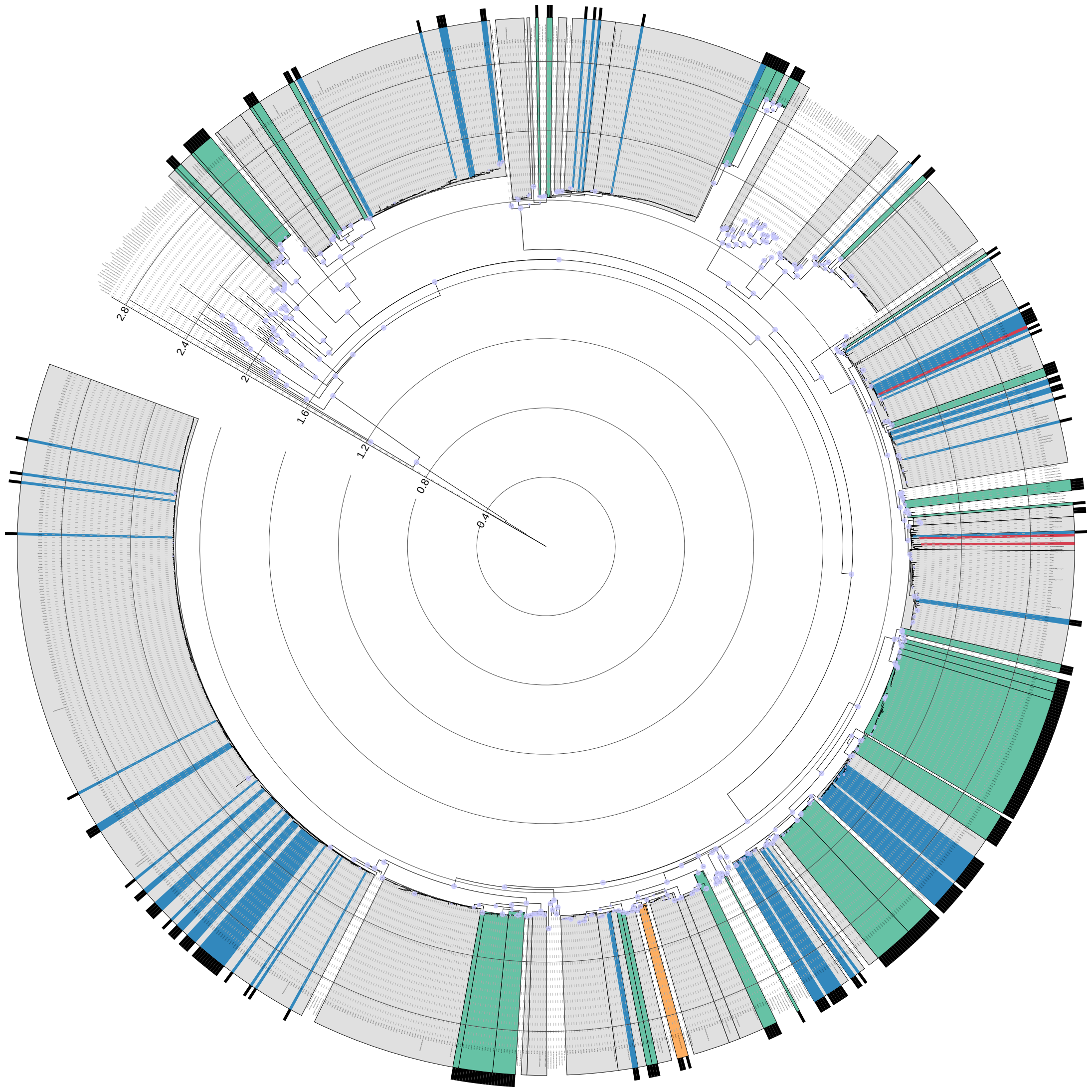
