## Supplementary File S3 for "Exploring eco-evolutionary and temporal patterns of arbuscular mycorrhizal fungal communities colonizing *Sorghum bicolor* across sites of contrasting land use history and climate"

### Supplementary Figures

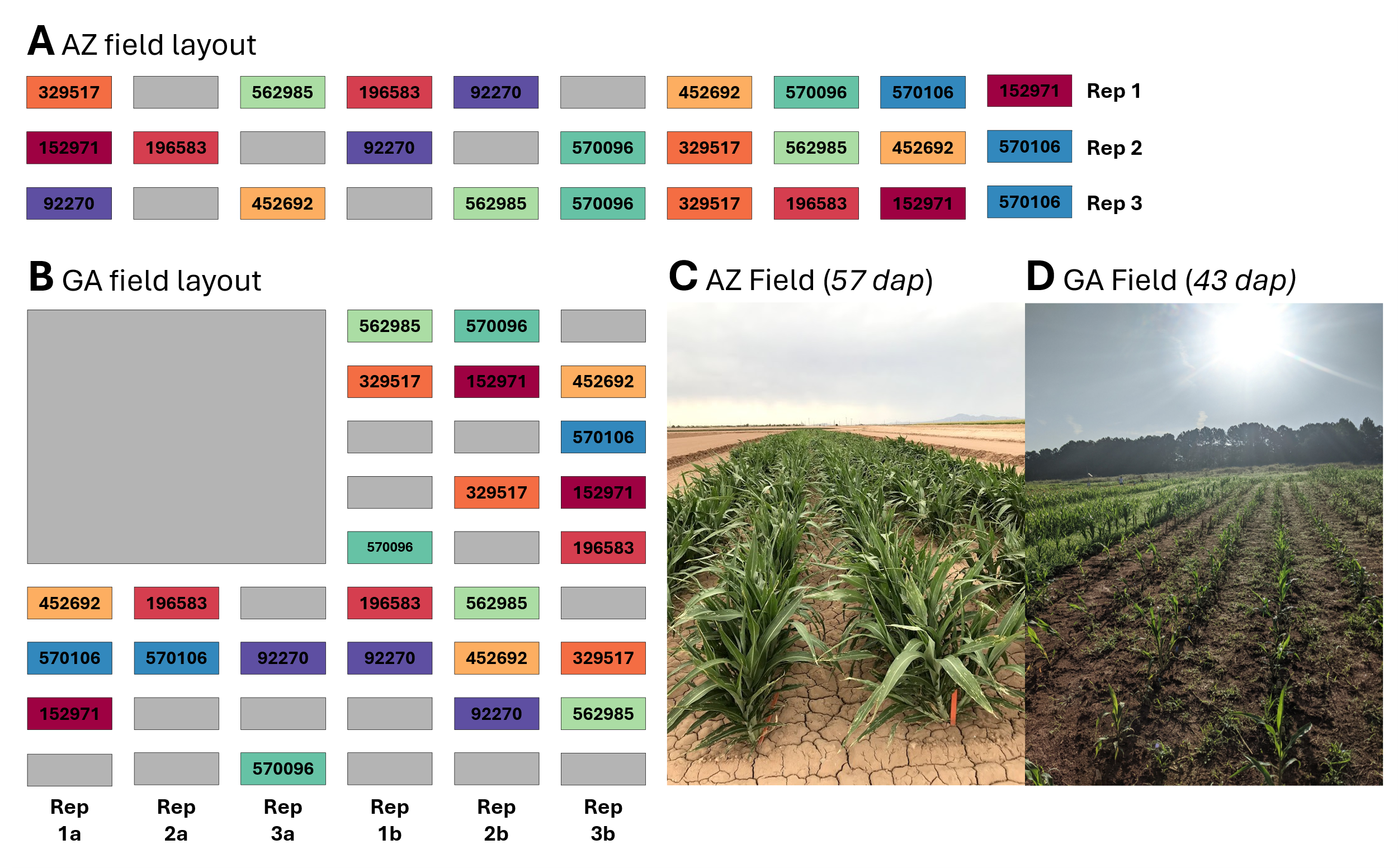

**Supplementary Figure S1.** Field Plans for **A)** AZ, and **B)** GA, with genotype plots colored by genotype. Plots were composed of single rows of 10 plants each, sowed 10cm apart from one another. Colored blocks represent genotype rows, which were composed of ten plants each spaced ~ 20 cm apart. Grey blocks are non-experimental plots. Plots were ~ 140 cm apart from one another within a row (displayed in horizontal orientation in this figure), and ~ 91 cm apart between rows. Demonstrative images are shown of **C** the AZ field site (taken June 23rd, 57 dap, TP3), and **D** the GA field site (taken July 14th, 43 days after planting, prior to collected timepoints).

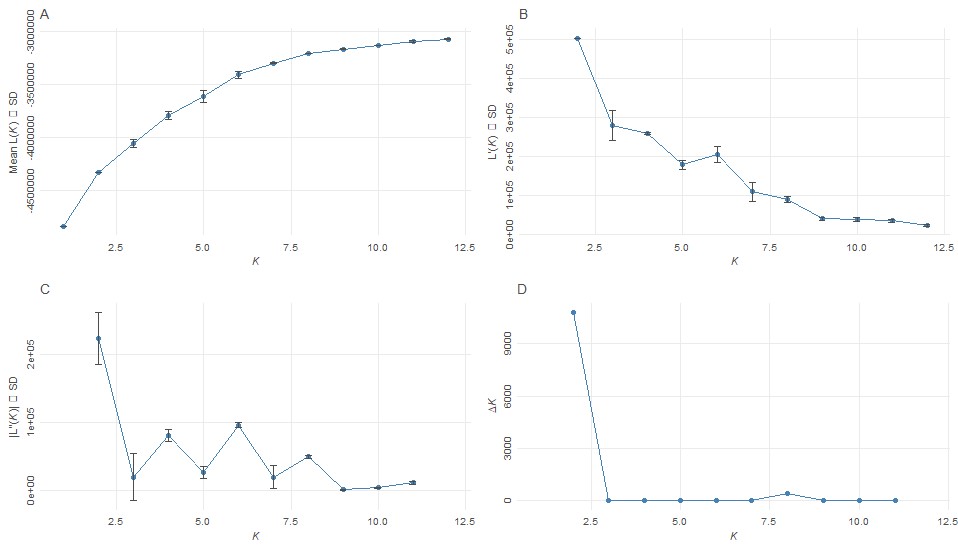

**Supplementary Figure S2.** Results of implementing the Evanno method to determine the optimum number of genetic clusters using the STRUCTURE analysis data of (Brenton et al. 2016) for the BAP panel of sorghum.

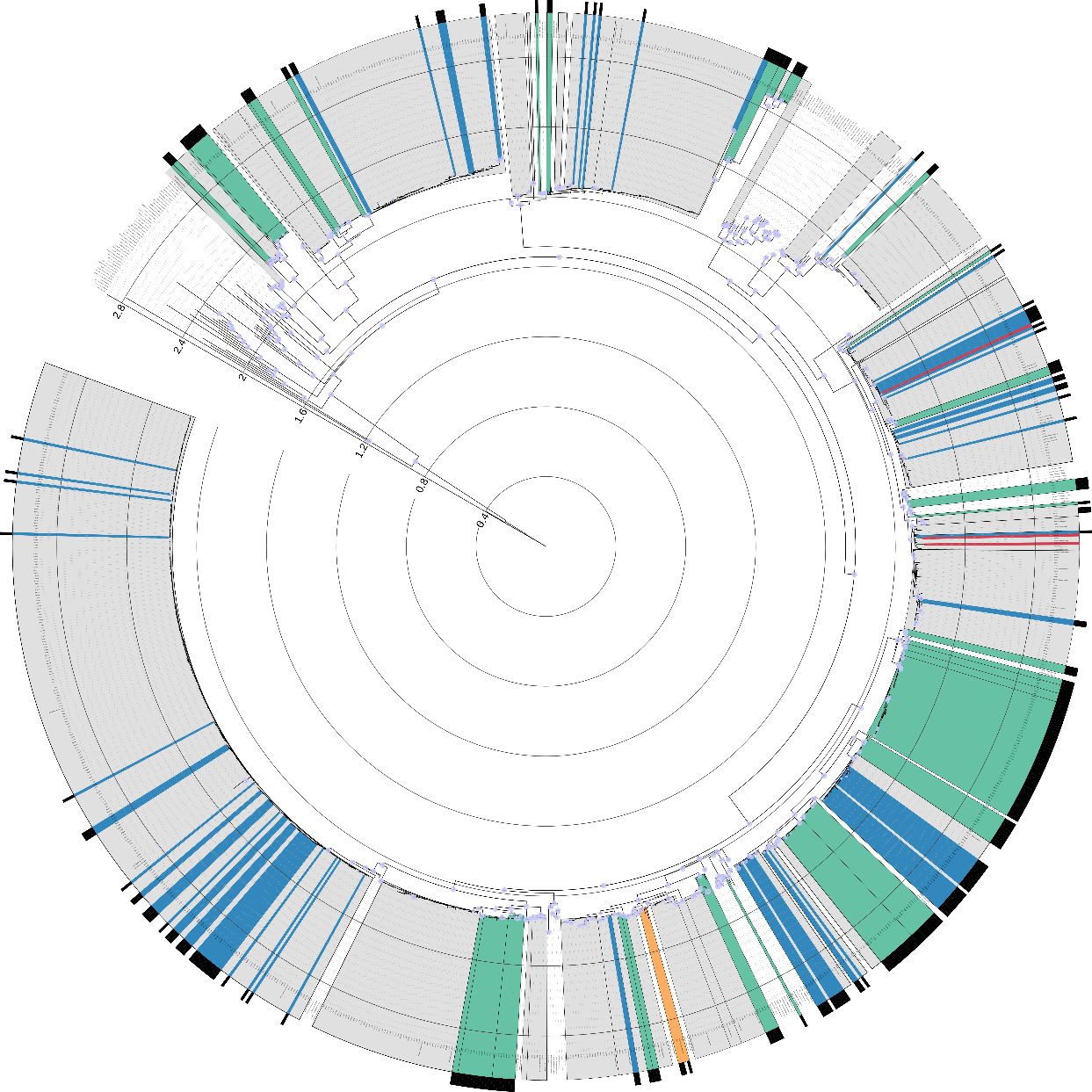

**Supplementary Figure S3.** Phylogeny of ASV sequences decorated upon the Delevaux reference backbone phylogeny (VS16_LSUDB_2024, (Delavaux et al., 2024) for AMF. Only bootstrap values > 50 are shown as blue circles on the appropriate nodes. Phylogenetic distance is shown nested within the tree. One backbone sequence (*Silvaspora neocaledonica*) was removed from the visualization due to having a suspiciously long branch length relative to other entries. Grey boxes aggregate species-level assignments made through the QIIME2 naïve Bayesian feature classification model trained on the EUKARYOME LSU database. Black bars at the periphery of the phylogeny denote ASVs that were not assigned to a species-level classification through the QIIME2 classifier. Green boxes aggregate where we manually defined putative novel molecular taxa (MTs) based on a mixture of branch support (> 50%) and clear distinction of phylogenetic distance from other closely related sequences. Blue denotes ASVs that were manually re-annotated to a species that had previously been defined through feature classification but was not assigned to the ASV in this first pass. Yellow denotes ASVs that were assigned to the nearest labeled sequence within the phylogeny based on the Delavaux taxonomy but was not represented in any EUKARYOME assignments. Within the grey boxes, red denotes where existing annotation from the feature classifier was manually re-annotated. This includes two ASVs assigned to *Gigaspora rosea* that clustered within the *Gigaspora gigantea* clade, and one ASV that was assigned to *Racocetra* but clustered within the closely related *Cetraspora* genus. See Supplementary File S2 for the high-resolution searchable PDF of the phylogeny.

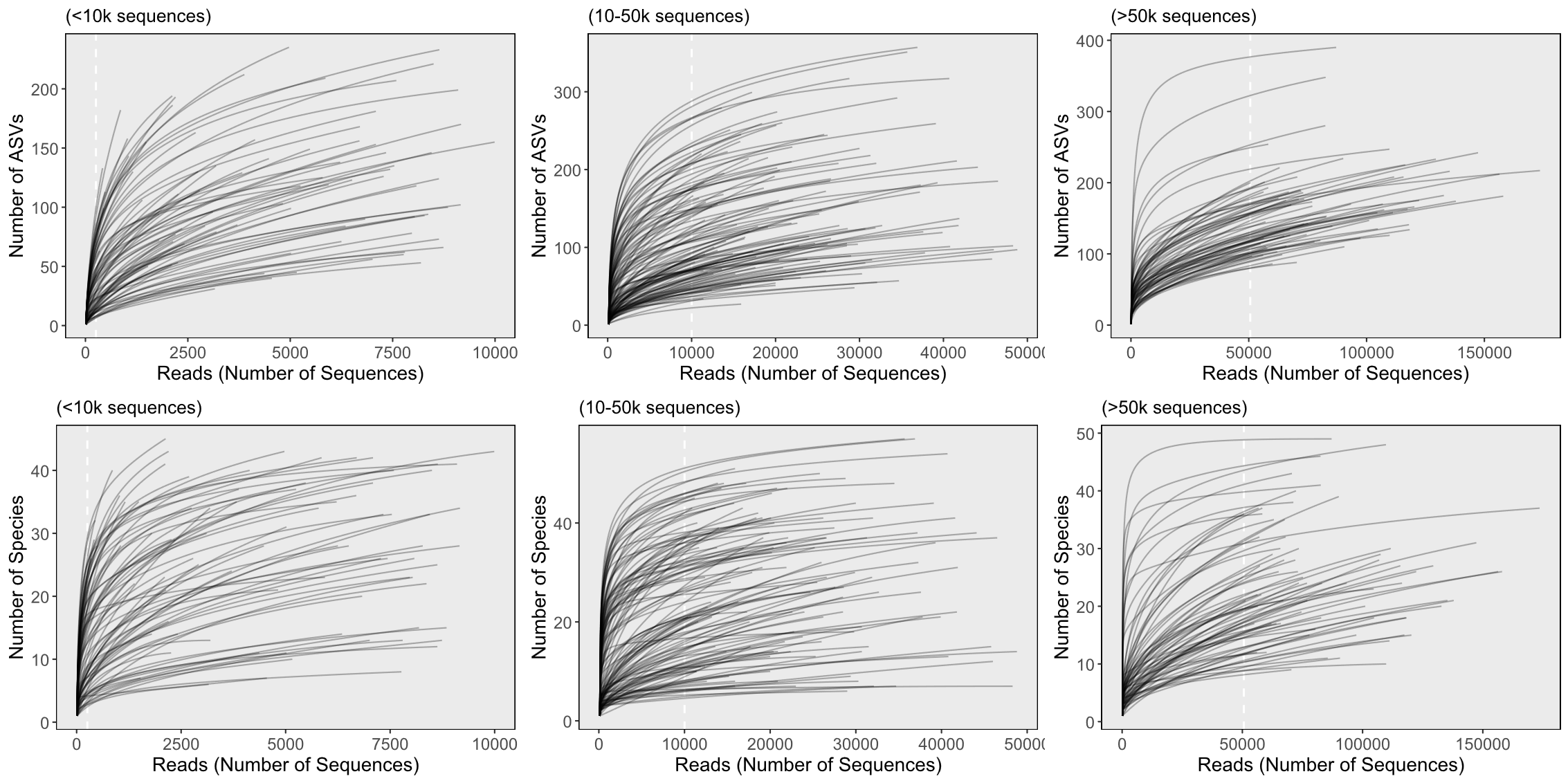

**Supplementary Figure S4.** Rarefaction curves generated based on ASV and aggregated species richness per sample for samples with depths between 0-10k, between 10k-50k, and over 50k. Sequencing depths refer to the number of AMF sequences obtained per sample, not the total sequencing depth.

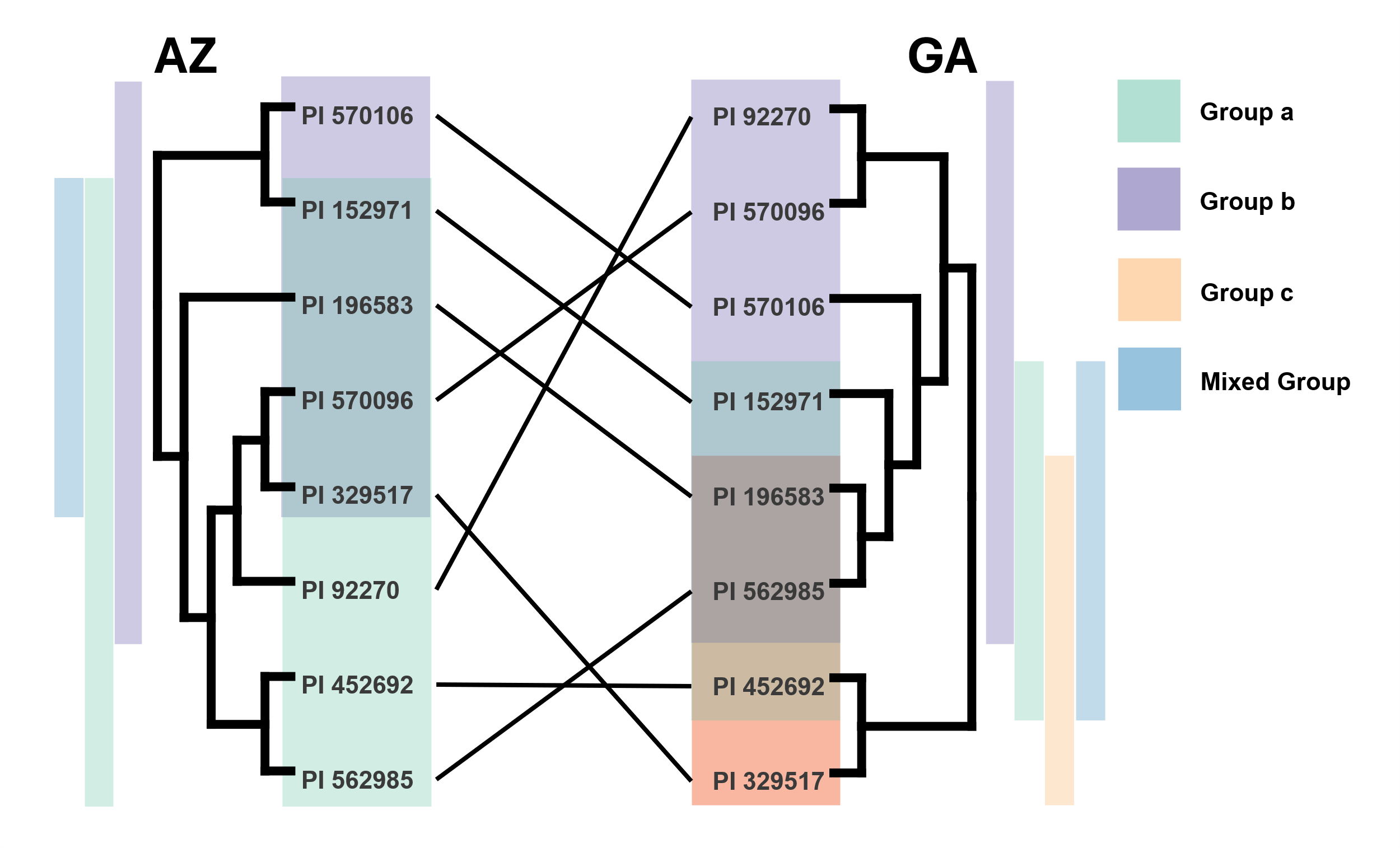

**Supplementary Figure S 5.** Tanglegram of dendrograms constructed from hierarchical clustering of Bray-Curtis distances calculated from the average AMF communities of each sorghum genotype per site. Colors represent groupings of statistical similarity (BH-adjusted p < 0.05) after pairwise PERMANOVA testing of between-genotype differences within each site. Group labels a and b to arbitrary simply to denote clusters of genotypes with highly divergent AMF communities.

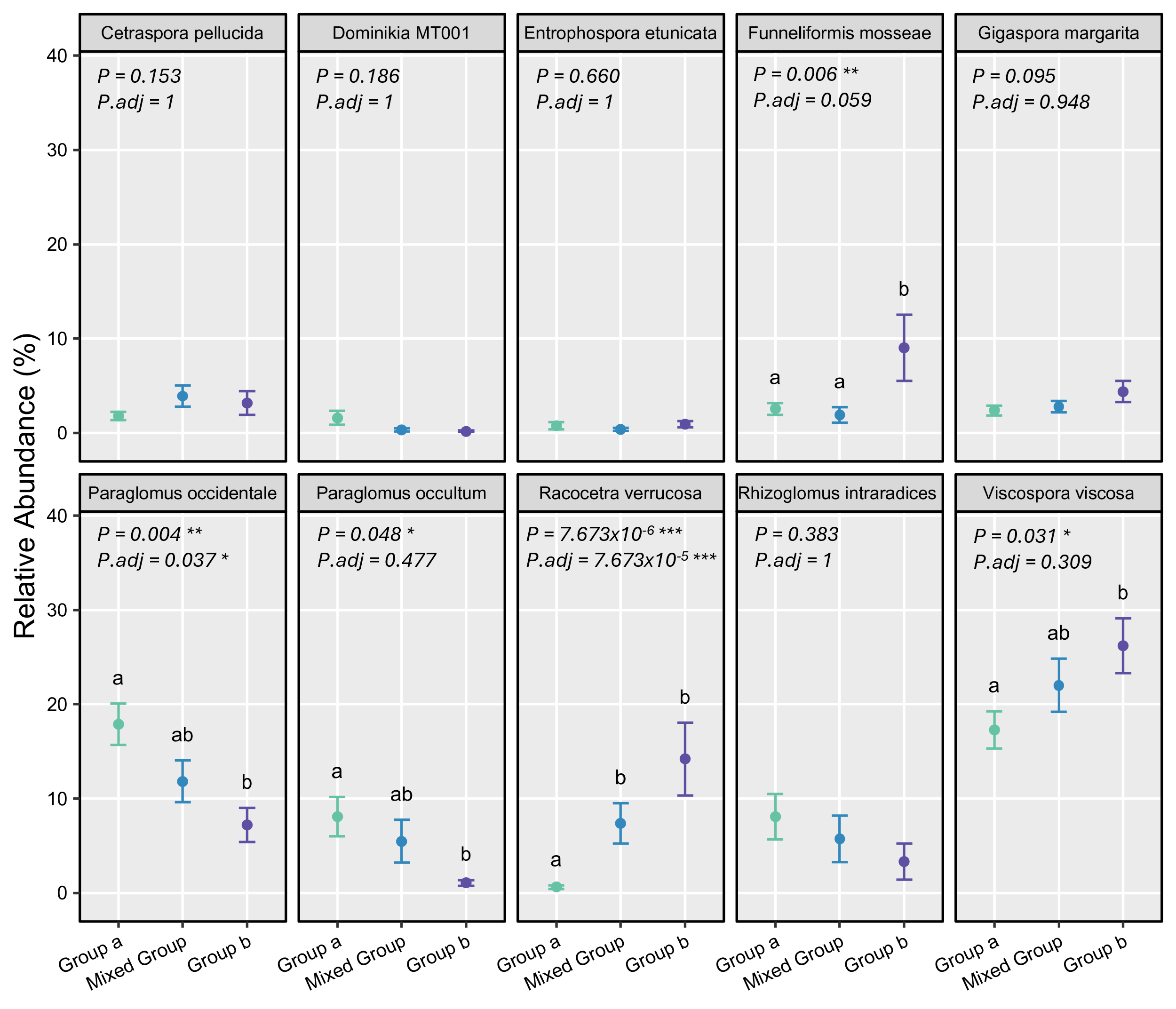

**Supplementary Figure S6.** Mean relative abundance ±SE of then 10 AMF species / molecular taxa with the highest contribution to the constrained CAP ordination of AMF for AZ. P-values from ANOVA testing of linear mixed effect models are shown. Both uncorrected and corrected (BH) p-values are shown. Letters represent groupings of statistical similarity (p < 0.05) based on pairwise Tukey post-hoc testing between Genotype groups. Genotype groups for analysis were defined based on whole community PERMANOVA analyses. Group a = PI 452692, PI 562985, PI 92270; Group b = PI 570106; Mixed Group = PI 152971, PI 196583, PI 329517, PI 570096. Replication: n=14 for PI 196583 and PI 452692, n=15 for all other genotypes.

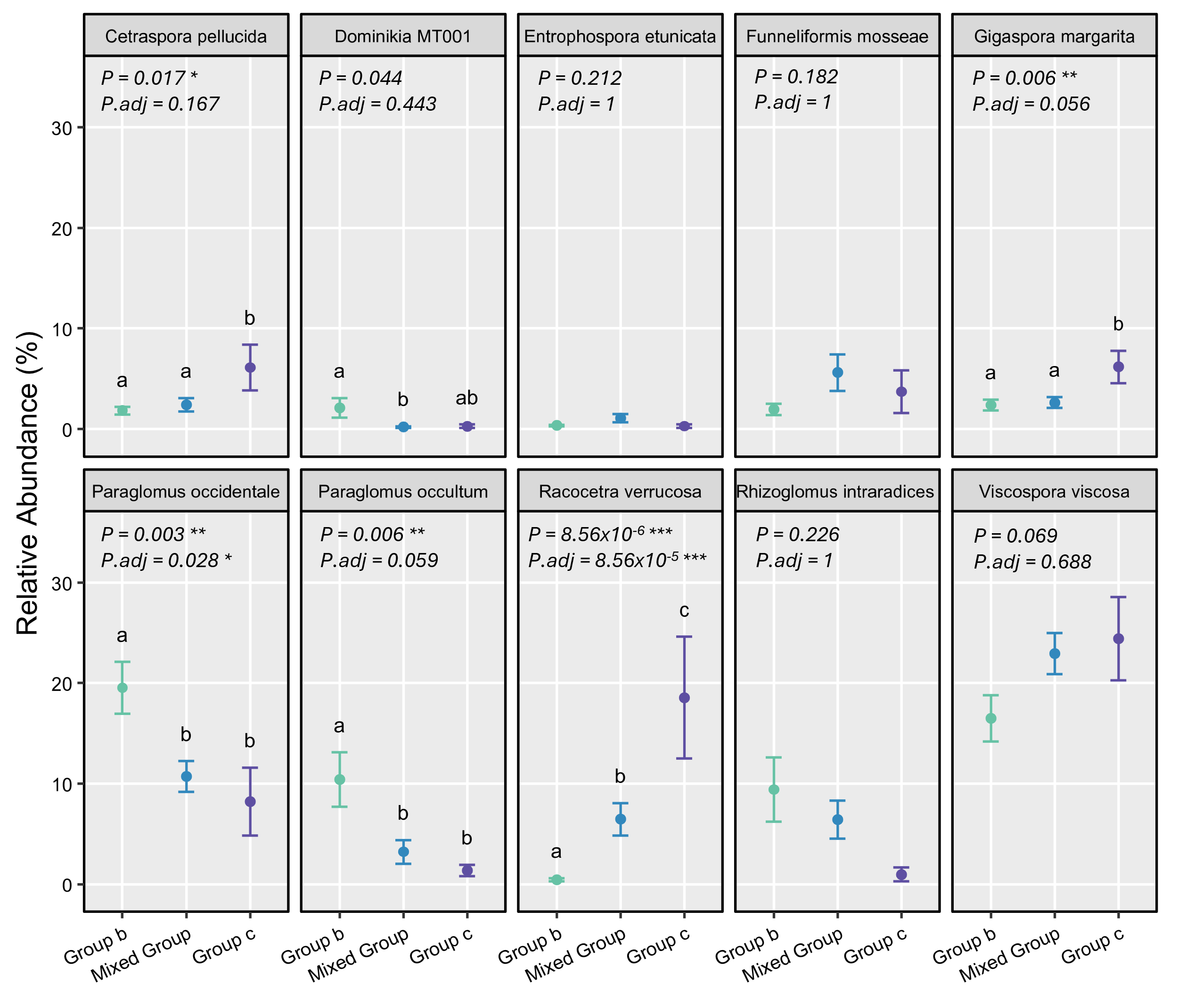

**Supplementary Figure S7.** Mean relative abundance ±SE of then 10 AMF species / molecular taxa with the highest contribution to the constrained CAP ordination of AMF for GA. P-values from ANOVA testing of linear mixed effect models are shown. Both uncorrected and corrected (BH) p-values are shown. Letters represent groupings of statistical similarity (p < 0.05) based on pairwise Tukey post-hoc testing between Genotype groups. Group a = PI 157971, PI 570096, PI 570106, PI 92270; Group b = PI 329517, PI 562985; Mixed Group = PI 196583, PI 562985. Replication: n=14 for PI 196583 and PI 452692, n=15 for all other genotypes.

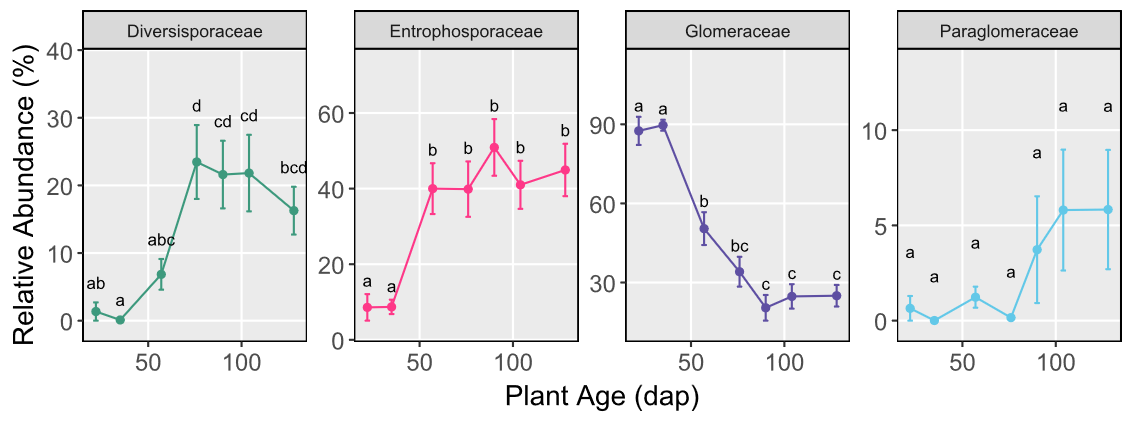

**Supplementary Figure S8**. Mean relative abundances ± SE of AMF families with significantly different (p < 0.05) abundances across timepoints in AZ, tested through general additive mixed models (GAMM). Letters represent groupings of statistical similarity (fdr-adjusted p < 0.05) after post-hoc pairwise testing following linear mixed effect (lme) models. Replication: n = 22 for TP4 / 76 dap. n= 24 for all other timepoint / plant age classes.

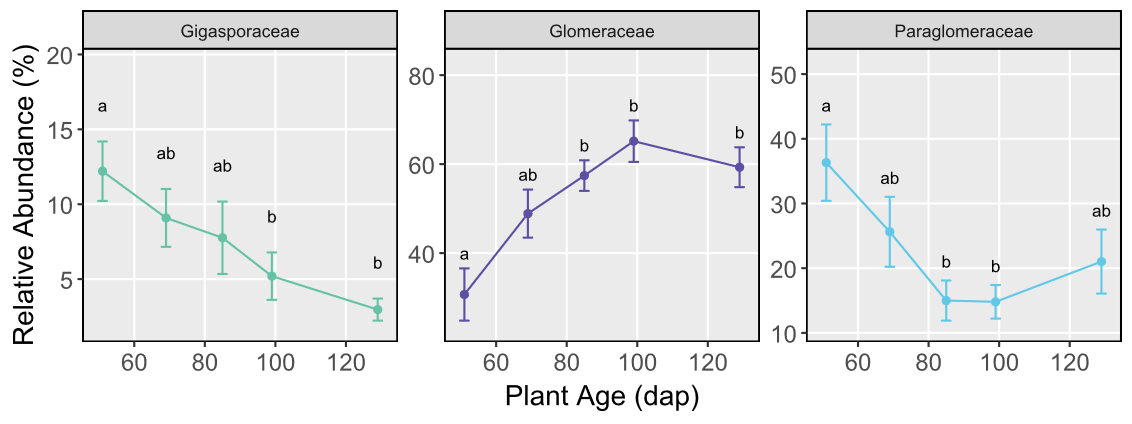

**Supplementary Figure S9.** Mean relative abundances ± SE of AMF families with significantly different (p < 0.05) abundances across timepoints in GA, tested through general additive mixed models (GAMM). Letters represent groupings of statistical similarity (fdr-adjusted p < 0.05) after post-hoc pairwise testing following linear mixed effect (lme) models. Replication: n = 24 per timepoint / plant age.

### Supplementary Tables

**Supplementary Table S1**. Genotypes used in this study with their characteristics and STRUCTURE-defined genetic sub-populations and phenotypes per (Brenton et al., 2016).

| Genotype | Name / Cultivar | Type | Photoperiod | K = 6 Population | K = 8 Population |
| --- | --- | --- | --- | --- | --- |
| PI 452692 | ETS 2110 | Cellulosic | Sensitive | 1 | 1 |
| PI 152971 | AWANLEK | Sweet | Insensitive | 2 | 2 |
| PI 196583 | MN 3080 | Sweet | Insensitive | 3 | 3 |
| PI 92270 | IS 12748 | Sweet | Insensitive | 4 | 4 |
| PI 570096 | IS 22874 | Cellulosic | Sensitive | 5 | 5 |
| PI 329517 | IS 11269 | Cellulosic | Sensitive | 5 | 6 |
| PI 562985 | AYI | Cellulosic | Sensitive | 5 | 7 |
| PI 570106 | IS 22884 | Cellulosic | Sensitive | 6 | 8 |

### Supplementary Results

#### Results S1: Aboveground plant characteristics

Sorghum performed better in AZ, exhibiting on average ~100**%** higher mean dry biomass at the final harvest (TP7) in AZ than GA. There was not a statistically significant GxE contribution to biomass (Supplementary Figure S9 A), and genotypes generally exhibited a strong correlation (R^2^ = 0.749, p = 0.033) of biomass phenotypes between sites (Supplementary Figure S10 A). Mean height was similar between sites (Mean height AZ: 2.54 m; GA: 2.68 m), though there was a significant GxE interaction for this trait at both locations. While the overall ranking of height phenotypes mostly agreed between sites, AZ showed a more distinct separation between genotypes (Supplementary Figure S9 B). Mean height values show a weaker relationship between sites, having a high correlation value but no statistical significance (R^2^ = 0.607, p = 0.111,Supplementary Figure S10 B).

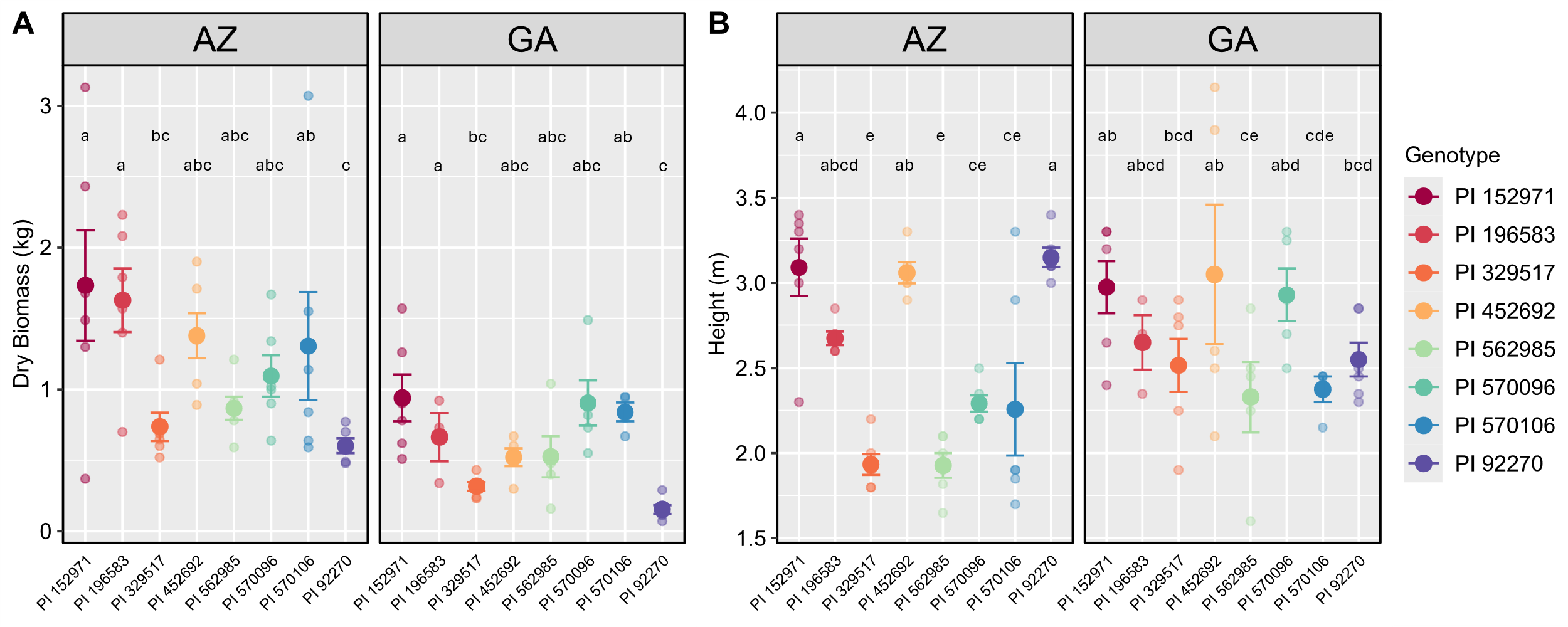

**Supplementary Figure S10.** Mean ± SE values of **A)** aboveground dry plant biomass (kg) and **B)** plant height (m), separated by sorghum genotype and site. Letters represent groupings of statistical similarity (fdr-adjusted p < 0.05) after pairwise testing based on **A)** between-genotype comparisons and **B)** genotype x environment interactions, chosen through global linear model testing. Replication: AZ, n=5 for PI 452692 (height only) and PI 92270 (biomass only), n=6 for all other genotypes for both variables; GA, n=6 per genotype.

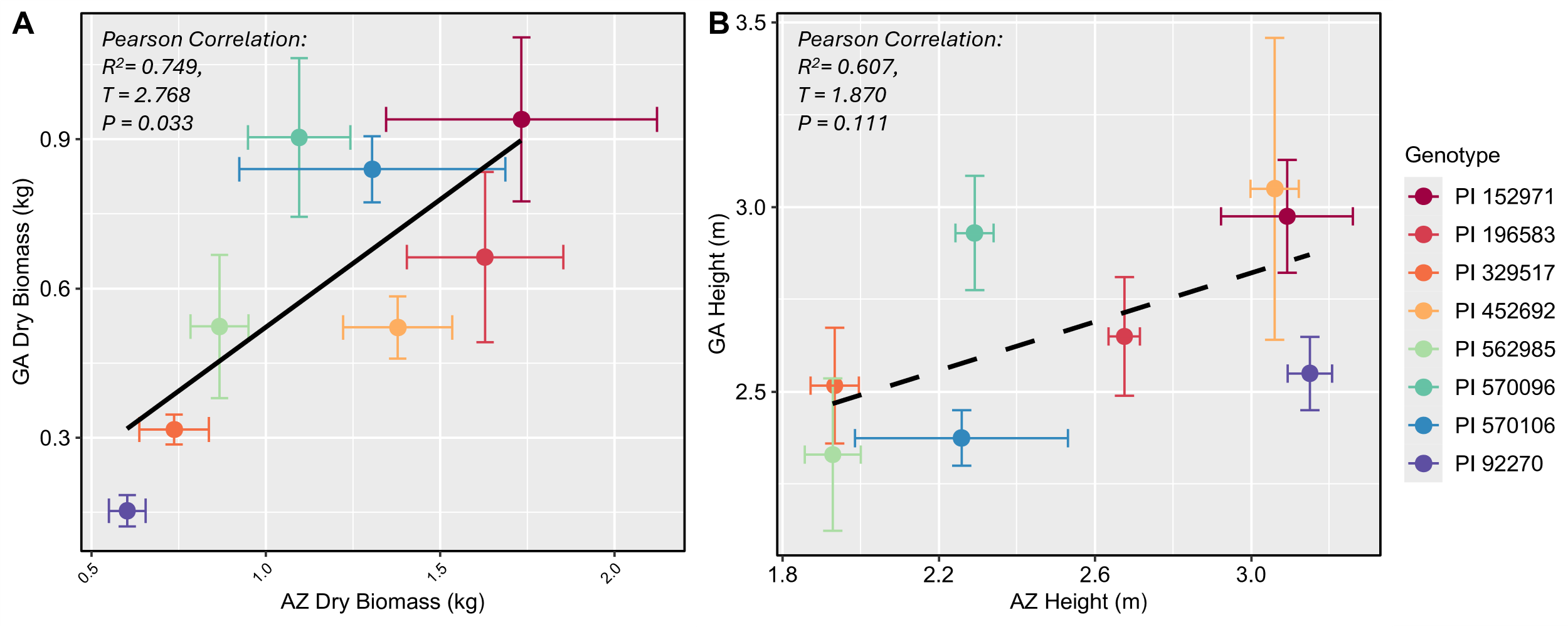

**Supplementary Figure S11.** Correlations between mean **A)** dry biomass (kg) and **B)** height (m) values per genotype in AZ and GA at TP7. Bars represent standard error of the mean value per genotype at both sites. Replication: AZ, n=5 for PI 452692 (height only) and PI 92270 (biomass only), n=6 for all other genotypes for both variables; GA, n=6 per genotype. Correlation analysis was performed only using mean values and not the individual plant values as samples are not explicitly paired between sites.

Brenton, Z. W., Cooper, E. A., Myers, M. T., Boyles, R. E., Shakoor, N., Zielinski, K. J., Rauh, B. L., Bridges, W. C., Morris, G. P., & Kresovich, S. (2016). A Genomic Resource for the Development, Improvement, and Exploitation of Sorghum for Bioenergy. *Genetics*, *204*(1), 21–33. https://doi.org/10.1534/genetics.115.183947

Delavaux, C. S., Ramos, R. J., Stürmer, S. L., & Bever, J. D. (2024). An updated LSU database and pipeline for environmental DNA identification of arbuscular mycorrhizal fungi. *Mycorrhiza*, *34*(4), 369–373. https://doi.org/10.1007/s00572-024-01159-3
